# Human pancreatic islet microRNAs implicated in diabetes and related traits by large-scale genetic analysis

**DOI:** 10.1101/2022.04.21.489048

**Authors:** Henry J. Taylor, Yu-Han Hung, Narisu Narisu, Michael R. Erdos, Matthew Kanke, Tingfen Yan, Caleb M. Grenko, Amy J. Swift, Lori L. Bonnycastle, Praveen Sethupathy, Francis S. Collins, D. Leland Taylor

## Abstract

Genetic studies have identified ≥240 loci associated with risk of type 2 diabetes (T2D), yet most of these loci lie in non-coding regions, masking the underlying molecular mechanisms. Recent studies investigating mRNA expression in human pancreatic islets have yielded important insights into the molecular drivers of normal islet function and T2D pathophysiology. However, similar studies investigating microRNA (miRNA) expression remain limited. Here, we present data from 63 individuals, representing the largest sequencing-based analysis of miRNA expression in human islets to date. We characterize the genetic regulation of miRNA expression by decomposing the expression of highly heritable miRNAs into *cis-* and *trans-*acting genetic components and mapping *cis*-acting loci associated with miRNA expression (miRNA-eQTLs). We find (i) 81 heritable miRNAs, primarily regulated by *trans*-acting genetic effects, and (ii) 5 miRNA-eQTLs. We also use several different strategies to identify T2D-associated miRNAs. First, we colocalize miRNA-eQTLs with genetic loci associated with T2D and multiple glycemic traits, identifying one miRNA, miR-1908, that shares genetic signals for blood glucose and glycated hemoglobin (HbA1c). Next, we intersect miRNA seed regions and predicted target sites with credible set SNPs associated with T2D and glycemic traits and find 32 miRNAs that may have altered binding and function due to disrupted seed regions. Finally, we perform differential expression analysis and identify 13 miRNAs associated with T2D status—including miR-187-3p, miR-21-5p, miR-668, and miR-199b-5p—and 4 miRNAs associated with a polygenic score for HbA1c levels—miR-216a, miR-25, miR-30a-3p, and miR-30a-5p.

## Introduction

Type 2 diabetes (T2D) is a leading contributor to global morbidity and mortality (1) and is characterized by reduced insulin response in insulin-sensitive tissues and pancreatic islet beta cell dysfunction (2, 3). However, despite our knowledge of the role of insulin in the disease, the underlying molecular and cellular mechanisms driving T2D are not wholly understood. Recent studies investigating circulating biomarkers of T2D identified several microRNAs (miRNAs) with altered levels in diabetic patients (4–9), raising the possibility that miRNAs play an important role in human T2D pathophysiology.

Mature miRNAs are small (∼22 nucleotides long), non-coding RNA molecules that typically function as post-transcriptional gene regulators by tethering the miRNA-induced silencing complex (miRISC) to target mRNAs, leading to mRNA silencing and degradation (10–12). Several seminal studies have implicated miRNAs in pancreatic islet development and function (13–24). However, the bulk of these studies are in model systems (e.g., animal models, cell lines), so our knowledge of miRNAs in human pancreatic islets (HPIs) remains limited. To date, most studies examining miRNA expression in HPIs have tested only a small number of miRNAs (25–28). Of the two more comprehensive sequencing-based studies of HPI miRNAs (29, 30), both have been limited in sample size (n≤7).

In addition, incorporating genetic information in the analysis of miRNA expression in HPIs may help identify miRNAs with a causal role in T2D pathophysiology. To date, ≥240 loci have been reported to be associated with T2D risk, yet the majority of these signals lie in non-coding regions of the genome, masking the underlying molecular mechanisms (31, 32). Previous studies have successfully integrated genetic and RNA-seq data in HPIs to characterize the genetic drivers of mRNA expression and nominate candidate causal genes for T2D by identifying shared genetic signals between loci associated with islet gene expression (eQTLs) and T2D risk (33–37). However, such integrative analyses have yet to be performed for miRNAs in HPIs.

Here, we present data from 63 individuals, representing the largest sequencing-based analysis of miRNA expression in HPIs to date (Fig. 1; Table S1; 57 with genotypes). Using genotype information (n=57), we estimate the SNP-based heritability for each miRNA (i.e., the proportion of variation in expression that can be explained by additive effects of common genetic variants) and find 81 highly heritable miRNAs. To perform a comparative analysis of heritability between miRNAs and mRNAs, we also estimate the SNP-based heritability of mRNAs using a subset of islets (n=40) with genotype and mRNA expression data. We show that regulation of heritable miRNAs is primarily driven by *trans*-acting genetic effects, whereas heritable mRNAs are regulated by a combination of *cis-*acting and *trans-*acting genetic effects. Given the importance of islet dysfunction in T2D pathophysiology, we apply several different strategies to assess the role of miRNA expression in diabetes. First, we map miRNA-eQTLs and colocalize the 5 identified miRNA-eQTLs with genetic signals from association studies of T2D and glycemic traits. Next, we identify SNPs from these genetic studies that are in the 99% credible sets (i.e., variants that are most likely to be the causal variant at each independent genetic association signal) and lie in miRNA seed regions or predicted target sites. Finally, using all miRNA expression data (n=63), we identify miRNAs associated with T2D status and polygenic scores (PGSs) of T2D-related traits, as well as sex, age, and body mass index (BMI).

**Fig. 1.**
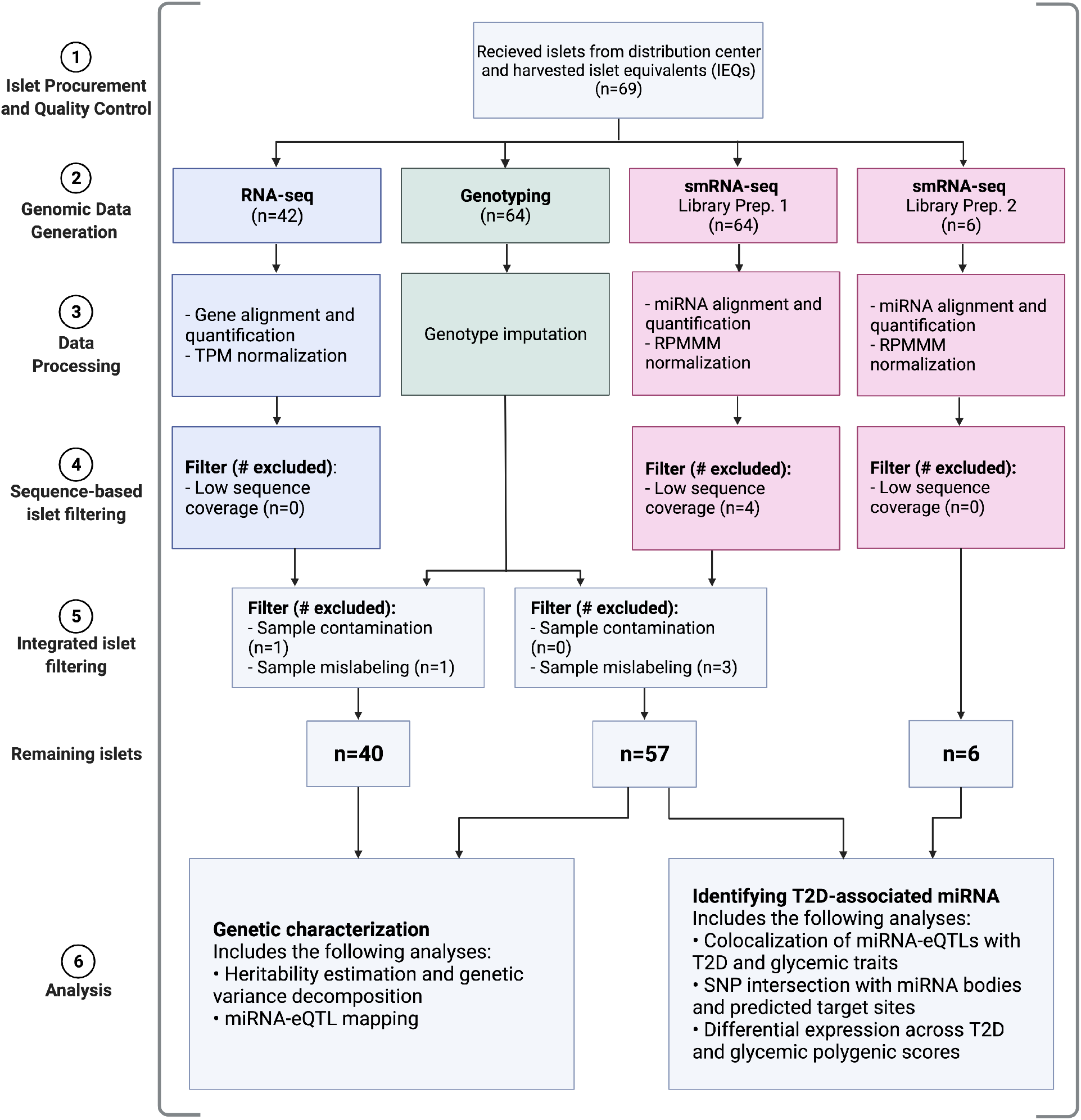
Study overview. Colors correspond to data modalities. Created with BioRender.com.

## Results

### Data overview

We procured 69 human pancreatic islet (HPI) samples and performed small RNA-sequencing (smRNA-seq) on 63 samples, RNA-sequencing (RNA-seq) on 40 samples, and genotyping on 57 samples (Fig. 1; Fig. S1; Table S1). For the smRNA-seq, we generated data through two experiments: library preparation 1 (LP1; 57 samples) and library preparation 2 (LP2; 6 samples). In LP1, we generated an average of 38.87 million read pairs per sample (±14.45 standard deviation (SD); minimum read count ≥19.98 million), with a mean read length of 23.24 nucleotides (Fig. S2A). In LP2, we generated an average of 64.36 million read pairs per sample (±4.18 SD; minimum read count ≥58.61 million), with a mean read length of 22.62 nucleotides (Fig. S2B). Across both library preparations, we found that the microRNAs (miRNAs) identified by previous mouse and human studies (17, 29, 38) to be most abundant in islets (e.g., miR-375) were also the most abundant miRNAs in the data generated in this study (Fig. S3). In total, we identified 2,959 unique miRNAs, including 1,989 miRNA isoforms (isomiRs). Of these, 2,279 are either canonical (reference) miRNAs or isomiRs with nucleotide shifts ≤2 in either direction, which we referred to as “high confidence”. For the RNA-seq, we generated an average of 57.23 million read pairs per sample (±16.36 SD; minimum read count ≥21.49 million). Similar to smRNA-seq, we found that highly abundant mRNAs (e.g., *PRSS1, REG1A*) from previous HPI studies (35, 37) were also highly abundant in the islet mRNA data generated in this study (Fig. S4), underscoring the data quality.

### Heritability and selection pressure of miRNAs

Studies in mice hepatic and lung tissues (39, 40) suggest that the genetic architecture of miRNA expression is primarily driven by *trans*-acting factors, much more than mRNA which has a substantial *cis*-acting genetic component; however, to date, a comparison of miRNA and mRNA regulatory trends has yet to be explored in human islets. To compare the trends in genetic regulation of miRNA and mRNA species, we estimated the SNP-based heritability (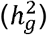) for miRNA and mRNA transcripts using imputed genotypes of common SNPs (Methods). We found that the fraction of heritable miRNA transcripts (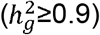) was substantially smaller than the fraction of heritable mRNA transcripts for both all miRNA transcripts (*P*=5.41×10^−5^, chi-squared test; Fig. 2A; Fig. S5A) and high confidence miRNA transcripts (P=1.09×10^−4^, chi-squared test; Fig. 2A; Fig. S5A), suggesting that miRNAs are under stronger selection pressure than mRNAs.

**Fig. 2.**
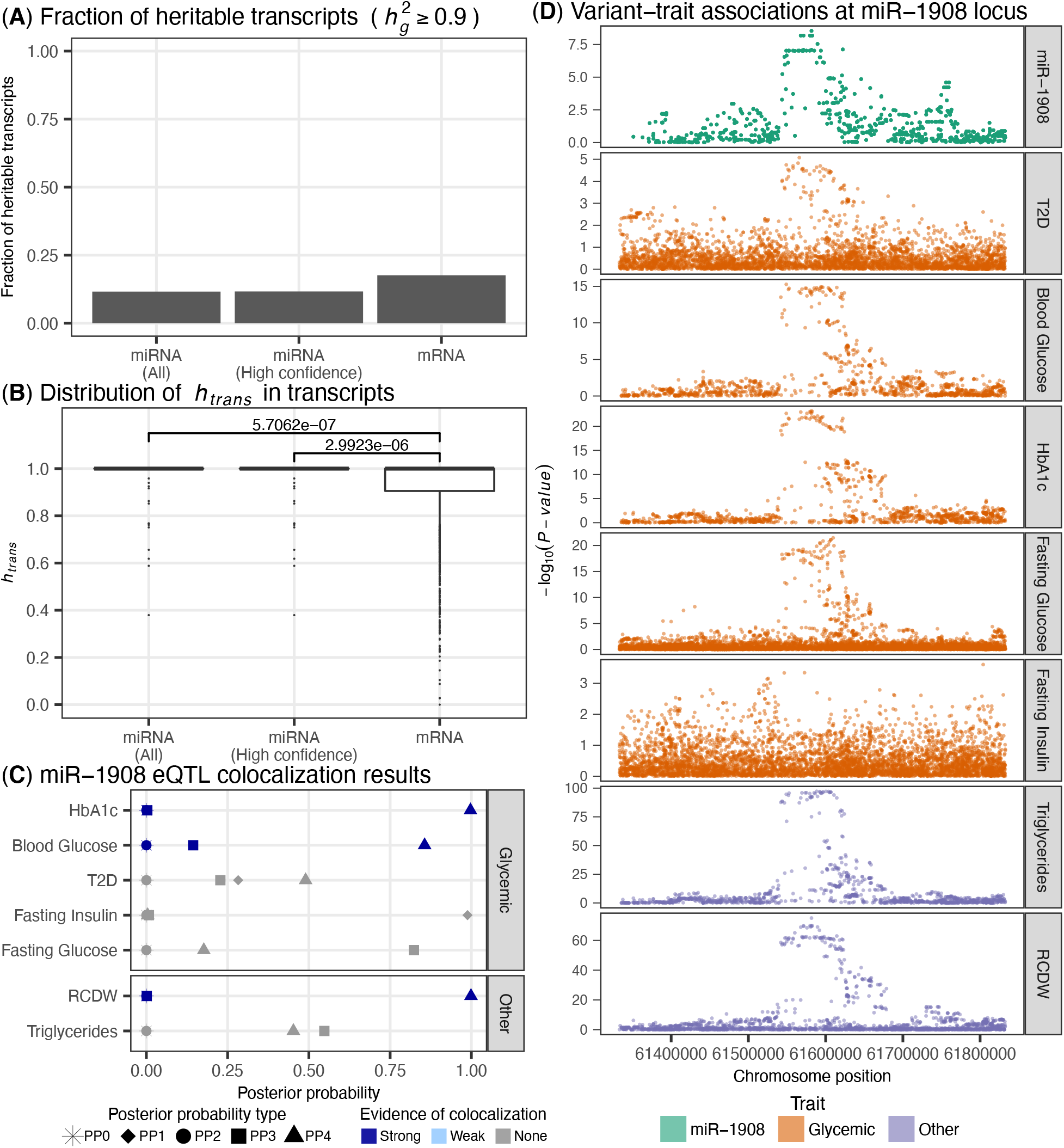
Genetic analysis of miRNA expression in human pancreatic islets. (A) Fraction of heritable transcripts (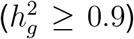) across miRNAs and mRNAs. (B) Distribution of variance explained by *trans*-acting genetic effects in expression of heritable transcripts. *P*-value calculated using Mann-Whitney rank test. (C) Evidence of genetic colocalization between miR-1908 eQTL and disease/trait genetic association signals (see Methods). Posterior probability definitions: PP0 - neither trait is associated, PP1 - miRNA-eQTL is associated, PP2 - GWAS is associated, PP3 - both traits are associated with different causal variants, PP4 - both traits are associated and share a single causal variant. We identified no traits showing weak evidence of colocalization. (D) For each disease/trait used in the colocalization analysis, Manhattan plots for all variants within a 250kb window on either side of the mature miR-1908 transcript.

We sought to understand the genetic architecture of heritable transcripts and decomposed the variance in transcript expression into *cis-* or *trans-*acting genetic effects, assuming that proximal SNPs (i.e., SNPs found within 250kb of the mature miRNA transcript or mRNA transcription start site) function primarily via *cis*-acting mechanisms and distal SNPs (i.e., outside of a 250kb window) function primarily via *trans*-acting mechanisms, as has been shown previously (41). We found that the variance explained by *trans*-effects (*h*_*trans*_) was greater in heritable miRNA transcripts than in heritable mRNA transcripts (*P*=5.71×10^−7^, Mann-Whitney rank test; Fig. 2B; Fig. S5B). These trends held true when just considering high confidence miRNA transcripts (*P*=2.99×10^−6^, Mann-Whitney rank test; Fig. 2B; Fig. S5B) as well as a larger 20Mb *cis* window size, to account for the fact that some miRNA transcription start sites (TSSs) may be quite far away from the miRNA body (*P*=8.53×10^−3^, Mann-Whitney rank test; Fig. S6) (42). Combined, these results suggest the genetic regulatory architecture of miRNAs differs from mRNAs as miRNA genetic regulation is primarily driven by *trans*-effects while mRNA is driven by a combination of *trans-* and *cis-*effects. Such a model is consistent with previous studies in rodents (39, 40).

### Identification of genetic effects on miRNA expression

We tested for genetic associations with miRNA expression in order to identify miRNA expression quantitative trait loci (miRNA-eQTLs; Methods). At a false discovery rate (FDR) of ≤5%, we found 5 miRNA-eQTLs (Table S3). We performed colocalization analysis between these miRNA-eQTLs and genetic loci associated with T2D and glycemic traits, including fasting blood glucose levels, blood glucose levels, fasting insulin levels, and glycated hemoglobin (HbA1c; Methods). We found no colocalizations with T2D. However, for one miRNA-eQTL, an eQTL for miR-1908 tagged by rs174559, we found strong evidence of colocalization with HbA1c and blood glucose levels (Fig. 2C-D). Since rs174559 is also strongly associated with triglycerides (TG) (43) and red blood cell distribution width (RCDW, a metric representing the heterogeneity of red blood cell volume) (44), we also performed colocalization with these traits. We found no evidence of colocalization with TG and strong evidence of colocalization with RCDW (Fig. 2C), possibly suggesting pleiotropic effects of this locus across tissues.

Focusing on the islet-relevant colocalization with glycemic traits, we looked for (i) *cis-*effects of rs174559 on the transcription of nearby protein coding genes in HPIs and (ii) *trans*-effects of rs174559 on protein coding genes throughout the genome to identify candidate target transcripts of miR-1908. We performed colocalization analysis between the miRNA-eQTL signal for miR-1908 and genetic associations for gene and exon expression identified in an islet eQTL study spanning 420 individuals (33). We found strong evidence of colocalization with several *FADS1* exons and with gene level *FADS1* expression (Fig. S8A). Although miR-1908 is located within a *FADS1* exon (there are several *FADS1* isoforms), we found the strongest colocalization was with an exon ∼4.3kb away from miR-1908. We performed a mediation analysis using 34 islet samples for which we had genotype, mRNA expression, and miRNA expression information (Methods). We found the genetic effects on miR-1908 and *FADS1* expression to be independent of each other (*P*=0.859, causal inference test; Fig. S8B). Next, we looked for *trans*-associations between rs174559 and protein coding genes in 40 samples from this study as *trans-*eQTL summary statistics from previous islet eQTL studies were not available (Methods). We found no associations, which may be driven by no signal or the lack of power to detect *trans*-eQTL effects at this sample size.

Previous studies have mapped miRNA-eQTLs in blood (45, 46). To understand the tissue specificity of miRNAs genetic effects, we compared the effect sizes of SNP-miRNA pairs from HPI miRNA-eQTLs to blood miRNA-eQTLs (Methods). We found that although some effects are replicated across datasets (e.g., the HPI rs174559 miR-1908 eQTL is also found in Sonehara et al. (46)), the HPI effects were generally not correlated with the genetic effects observed in blood (Fig. S9; maximum Spearman’s rho=0.07), suggesting that genetic effects on miRNAs may to some extent be tissue/cell type specific.

Finally, we performed allelic imbalance analysis using the smRNA-seq reads (Methods) but did not identify any miRNAs that exhibited imbalance of transcribed alleles.

### Identification of T2D-related genetic effects on miRNA function

To identify genetic variants associated with T2D and T2D-related phenotypes that may affect islet miRNA function (as opposed to overall expression levels), we overlapped 99% credible set SNPs from genetic studies for T2D and other glycemic traits with (i) mature miRNA genomic coordinates (which includes areas outside of the seed region and may affect the ability of the miRNA to bind to targets) and (ii) genomic coordinates for predicted miRNA target regions (i.e., the miRNA binding site on the mRNA transcript; Methods; Table S4). We did not identify any SNPs within mature miRNA coordinates, but we identified 16 T2D, 8 HbA1c, and 1 blood glucose credible set SNPs within predicted miRNA target regions. Because SNPs within miRNA target regions would exert their effects on the target transcript via *cis*-acting mechanisms, we further explored the effects of these SNPs using an islet *cis*-eQTL study spanning 420 individuals (33). We found one SNP, rs1464569—in the 99% credible set for an HbA1c signal—that lies within *NICN1* at a miR-532-3p target site and is in high linkage disequilibrium (1000GENOMES phase_3 FIN=1.00, CEU=1.00, GBR=1.00) with the tag SNP, rs4955440, for the gene-level *NICN1* eQTL. To explore if rs1464569 functions as an eQTL for *NICN1* by perturbing miRNA binding, we tested for an interaction effect by modeling *NICN1* expression with rs1464569 and miR-532-3p expression using 34 samples with mRNA expression, miRNA expression, and genetic data (Methods). We found no evidence of an interaction effect (*P*>0.05), leaving the mechanism driving the genetic association at this locus unresolved.

### Differentially expressed miRNAs

Next, we sought to identify miRNAs associated with sex, age, BMI, T2D disease status, and PGSs for T2D and glycemic traits (Methods).

For sex, age, and BMI, we performed meta-analysis of differential expression results from this study (n=63 total from two cohorts) and a previous smRNA-seq study of HPIs (n=7) (29). We identified 1 BMI-associated and 22 sex-associated miRNAs (FDR≤5% and |log2(fold change (FC))|≥1; Fig. 3; Fig. S10,11). The large number of sex-associated miRNAs is consistent with a previous study describing substantial genomic differences between male and female HPIs (28). Notably, all the sex-associated miRNAs occurred on autosomes.

**Fig. 3.**
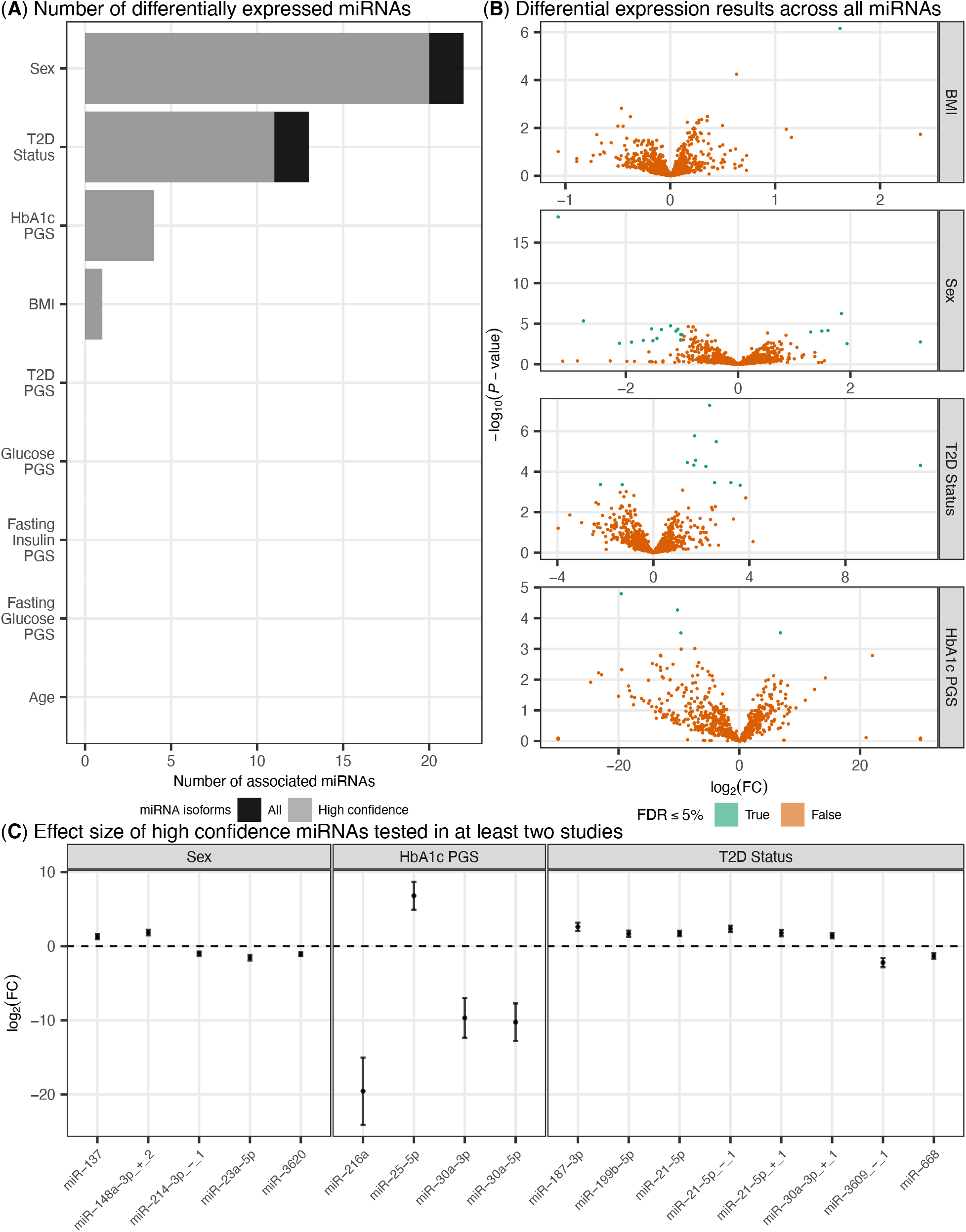
Differentially expressed miRNAs across diseases/traits. (A) Number of differentially expressed miRNAs (*log*_2_(*FC*) 1 and FDR 5%) across each disease/trait. (B) Volcano plots describing the *log*_2_(*FC*) (x-axis) and *log*_10_(*P*) (y-axis) of miRNAs across BMI, sex (male vs. female), T2D status, and HbA1c PGS. (C) Forest plot describing effect of high confidence differentially expressed miRNAs (*log*_2_(*FC*) 1 and FDR 5%). Effect sizes are from meta-analysis.

Similarly, for T2D status, we compared T2D (n=4) to normal glucose-tolerant (NGT, n=59) donors and meta-analyzed the differential expression results from this study with a previous study (29) consisting of 7 islet samples (n=4 T2D, n=3 NGT). We identified 13 miRNAs with altered expression in T2D (FDR≤5% and |log2(FC)|≥1), 12 of which were not identified previously in Kameswaran et al. (Fig. 3; Fig. S12), including 10 high confidence miRNAs such as miR-21-5p, miR-187-3p, miR-199b-5p, miR-668, and miR-4497.

For the polygenic scores, we used all miRNA samples with genotypes (n=57) and meta-analyzed differential expression results from individuals of European (n=42), African (n=7), and Hispanic/Latino (n=6) ancestries. We identified 4 miRNAs associated with the HbA1c PGS (FDR≤5% and |log2(FC)|≥1; Fig. 3; Fig. S13)—miR-216a, miR-25-5p, miR-30a-5p, and miR-30a-3p.

Finally, previous studies describe cell type heterogeneity as a possible confounder for bulk tissue gene expression studies (47, 48). To assess the contribution of cell type heterogeneity to the results described in this study, we estimated the fraction of alpha and beta cells, the primary cell types in processed HPIs (49–51), using miRNA profiles of sorted alpha and beta cells (Methods; Fig. S16) (29). For both the miRNA-eQTL (Fig. S17A) and differential expression analyses (Fig. S17B-E), we observed little change in the results of the models, suggesting cell type heterogeneity is not a primary driver of the results reported in this study.

## Discussion

MiRNAs are small, non-coding RNA molecules that serve as important post-transcriptional gene regulators in human health and disease (reviewed in Singh and Storey (52)). The critical contribution of miRNAs to proper pancreatic islet development and function as well as the dysregulation of miRNAs in the context of T2D is well documented by previous studies using rodent models and cell-based systems (13–24). Despite these advances, our knowledge of miRNA expression and activity in human islets remains limited.

In this study, we present the largest sequencing-based analysis of miRNA expression in HPIs to date (n=63). We describe the genetic regulation of HPI miRNA expression and provide insight into the role of miRNAs in T2D pathophysiology.

In the genetic analysis of HPIs, we show that expression of highly heritable miRNAs is driven primarily by *trans*-acting genetic effects, whereas expression of highly heritable mRNAs is driven by a combination of *cis-* and *trans-*acting genetic effects. This model is supported by previous studies comparing heritability of miRNAs and mRNAs in different tissues and models (39, 40) and suggests miRNAs may be under greater selective pressure than mRNAs, a hypothesis shared by a previous study investigating genetic variation in humans (53). Such selection pressures imply that to map miRNA-eQTLs thoroughly, large sample sizes will be required. Among the five identified miRNA-eQTLs, we find one miRNA-eQTL (miR-1908 tagged by rs174559) that colocalizes with genetic signals for multiple physiological traits—blood glucose, HbA1c, and RCDW—as well as gene- and exon-level expression of *FADS1*. However, we highlight two limitations. First, it is unclear whether these effects are mediated by miR-1908, *FADS1*, both, or neither. We perform a mediation analysis to disentangle directionality between the two transcripts, but we find the genetic effects on miR-1908 and *FADS1* to be independent. Second, it is unclear if the HbA1c and blood glucose effect is mediated by islets, blood, or some other tissue type. On the one hand, given (i) the rs174559 miR-1908 eQTL also occurs in blood (46), (ii) the RCDW effect at this locus, and (iii) extensive studies that document how erythrocytic (i.e., red blood cell related) genetic effects can mediate HbA1c levels (54–58), it is plausible that observed effect on HbA1c is mediated by blood. On the other hand, an erythrocytic mediated effect would not explain the observed blood glucose genetic effect. In addition, at this locus, a study partitioned HbA1c genetic associations into erythrocytic and glycemic effects (59). Using a SNP in high linkage disequilibrium with rs174559 (1000GENOMES phase_3 FIN=0.67, CEU=0.70, GBR=0.71) (60, 61), this study determined the effect at rs174559 on HbA1c was driven by glycemic effects rather than erythrocytic effects which would suggest the observed effects in islets may indeed be relevant.

In addition, we identify several HPI miRNAs differentially expressed across BMI, sex, T2D status, and HbA1c genetic scores. Of the miRNAs identified, 23 (60.52%) have been previously implicated at some level as being associated with a related phenotype (38, 62–76). Focusing on the miRNAs associated with T2D status and HbA1c genetic scores, these findings provide critical human islet-based confirmation of previous studies in rodents and cell lines. For example, we find HPI levels of miR-216a and miR-25 to be negatively and positively correlated with HbA1c genetic scores, respectively. In previous work, the miR-216a knockout mouse was reported to have defects in insulin secretion in the context of chronic high-fat diet (77), and miR-25 has been described as a negative regulator of insulin synthesis in INS-1 cells, a rat beta cell line (64). As another example, we show that miR-199b-5p is associated with T2D status, consistent with a previous study in murine beta cells where miR-199b-5p was shown to be highly responsive to glucose stimulation (78). We also find that miR-21-5p is strongly associated with T2D status. This observation comports with a recent study in which overexpression of miR-21-5p led to reduced glucose-stimulated insulin secretion in mice and human beta cells (79). Finally, we identify expression of miR-668, previously reported to be associated with T2D in human skeletal muscle (76), as a novel T2D association in HPIs.

Unraveling the genetic factors that contribute to pancreatic islet function and T2D has been a high priority for the diabetes research community. However, much of the focus has been on mRNA and chromatin. Here, we add data on another important source of functional variation. While this is the largest sequencing-based analysis of miRNA expression to date, even larger studies in the future may provide power to uncover more subtle contributions. Moreover, the application of single-cell approaches in the future may help to resolve miRNA differences in T2D across different islet cell types, and possibly even different subtypes of beta cells.

## Materials and Methods

A detailed description of computational and experimental analyses is provided in *Supplementary Materials and Methods*. Briefly, we performed small RNA sequencing (smRNA-seq) on 63 human pancreatic islet (HPI) samples and RNA sequencing (RNA-seq) on 40 HPIs—quantifying miRNA and mRNA expression. Using 63 HPIs with genotypes, we estimated the SNP-based heritability for miRNA (n=57) and mRNA (n=40) transcripts and decomposed the variance in expression of heritable transcripts into *cis-* and *trans*-acting genetic components using LIMIX v3.0.4. We tested for genetic variants associated with miRNA expression (miRNA-eQTLs) using LIMIX v3.0.4. To identify T2D-relevant miRNAs, we performed a colocalization analysis with miRNA-eQTLs and genetic loci associated with T2D and glycemic traits using coloc v3.1. To identify potential gene targets of miRNAs that colocalized with T2D or a glycemic trait, we also performed a colocalization analysis with miRNA-eQTLs and genetic loci associated with exon and gene expression in HPIs. We overlapped 99% credible set SNPs from genetic studies for T2D and glycemic traits with miRNA mature transcripts and target sites to identify variants that may alter miRNA function. Finally, we used DESeq2 v1.32.0 to identify miRNAs differentially expressed across T2D status, polygenic scores of T2D and glycemic traits, and other common phenotypes (i.e., sex, age, and BMI).

## Acknowledgements

This research was supported in part by United States’ National Institutes of Health (NIH) grants ZIA-HG000024 (to F.S.C.), Gates Cambridge Trust (to H.J.T.), the NIH Oxford-Cambridge scholars program (to H.J.T.), and American Diabetes Association (ADA) Pathway Award 1-16-ACE-47 (to P.S.). We thank Chad Krilow for supporting the work presented in this study.

## References

1. M. A. B. Khan, et al., Epidemiology of Type 2 Diabetes - Global Burden of Disease and Forecasted Trends. J. Epidemiol. Glob. Health 10, 107–111 (2020).

2. U. Galicia-Garcia, et al., Pathophysiology of type 2 diabetes mellitus. Int. J. Mol. Sci. 21 (2020).

3. S. E. Kahn, M. E. Cooper, S. Del Prato, Pathophysiology and treatment of type 2 diabetes: perspectives on the past, present, and future. Lancet 383, 1068–1083 (2014).

4. C. Wang, et al., Increased serum microRNAs are closely associated with the presence of microvascular complications in type 2 diabetes mellitus. Sci. Rep. 6, 20032 (2016).

5. Y. Rong, et al., Increased microRNA-146a levels in plasma of patients with newly diagnosed type 2 diabetes mellitus. PLoS ONE 8, e73272 (2013).

6. K. J. Belongie, et al., Identification of novel biomarkers to monitor β-cell function and enable early detection of type 2 diabetes risk. PLoS ONE 12, e0182932 (2017).

7. R. Jiménez-Lucena, et al., A plasma circulating miRNAs profile predicts type 2 diabetes mellitus and prediabetes: from the CORDIOPREV study. Exp. Mol. Med. 50, 1–12 (2018).

8. V. Ghai, et al., Circulating RNAs as predictive markers for the progression of type 2 diabetes. J. Cell. Mol. Med. 23, 2753–2768 (2019).

9. R. J. Farr, M. V. Joglekar, C. J. Taylor, A. A. Hardikar, Circulating non-coding RNAs as biomarkers of beta cell death in diabetes. Pediatr. Endocrinol. Rev. 11, 14–20 (2013).

10. J. O’Brien, H. Hayder, Y. Zayed, C. Peng, Overview of microrna biogenesis, mechanisms of actions, and circulation. Front Endocrinol (Lausanne) 9, 402 (2018).

11. P. J. Dexheimer, L. Cochella, Micrornas: from mechanism to organism. Front. Cell Dev. Biol. 8, 409 (2020).

12. D. P. Bartel, Metazoan MicroRNAs. Cell 173, 20–51 (2018).

13. M. N. Poy, et al., miR-375 maintains normal pancreatic alpha- and beta-cell mass. Proc Natl Acad Sci USA 106, 5813–5818 (2009).

14. M. Latreille, et al., MicroRNA-7a regulates pancreatic β cell function. J. Clin. Invest. 124, 2722–2735 (2014).

15. M. Kalis, et al., Beta-cell specific deletion of Dicer1 leads to defective insulin secretion and diabetes mellitus. PLoS ONE 6, e29166 (2011).

16. T. Melkman-Zehavi, et al., miRNAs control insulin content in pancreatic β-cells via downregulation of transcriptional repressors. EMBO J. 30, 835–845 (2011).

17. M. N. Poy, et al., A pancreatic islet-specific microRNA regulates insulin secretion. Nature 432, 226–230 (2004).

18. G. Xu, J. Chen, G. Jing, A. Shalev, Thioredoxin-interacting protein regulates insulin transcription through microRNA-204. Nat. Med. 19, 1141–1146 (2013).

19. B.-F. Belgardt, et al., The microRNA-200 family regulates pancreatic beta cell survival in type 2 diabetes. Nat. Med. 21, 619–627 (2015).

20. E. Roggli, et al., Changes in microRNA expression contribute to pancreatic β-cell dysfunction in prediabetic NOD mice. Diabetes 61, 1742–1751 (2012).

21. Q. Hu, et al., Obesity-Induced miR-455 Upregulation Promotes Adaptive Pancreatic β-cell Proliferation Through the CPEB1/CDKN1B Pathway. Diabetes (2022) https://doi.org/10.2337/db21-0134.

22. F. Zhang, et al., Obesity-induced overexpression of miR-802 impairs insulin transcription and secretion. Nat. Commun. 11, 1822 (2020).

23. M. V. Joglekar, V. M. Joglekar, A. A. Hardikar, Expression of islet-specific microRNAs during human pancreatic development. Gene Expr. Patterns 9, 109–113 (2009).

24. M. V. Joglekar, V. S. Parekh, S. Mehta, R. R. Bhonde, A. A. Hardikar, MicroRNA profiling of developing and regenerating pancreas reveal post-transcriptional regulation of neurogenin3. Dev. Biol. 311, 603–612 (2007).

25. D. Klein, et al., MicroRNA expression in alpha and beta cells of human pancreatic islets. PLoS ONE 8, e55064 (2013).

26. C. Bolmeson, et al., Differences in islet-enriched miRNAs in healthy and glucose intolerant human subjects. Biochem. Biophys. Res. Commun. 404, 16–22 (2011).

27. J. K. Ofori, et al., Human Islet MicroRNA-200c is Elevated in Type 2 Diabetes and Targets the Transcription Factor ETV5 to Reduce Insulin Secretion. Diabetes (2021) https://doi.org/10.2337/db21-0077.

28. E. Hall, et al., Sex differences in the genome-wide DNA methylation pattern and impact on gene expression, microRNA levels and insulin secretion in human pancreatic islets. Genome Biol. 15, 522 (2014).

29. V. Kameswaran, et al., Epigenetic regulation of the DLK1-MEG3 microRNA cluster in human type 2 diabetic islets. Cell Metab. 19, 135–145 (2014).

30. M. van de Bunt, et al., The miRNA profile of human pancreatic islets and beta-cells and relationship to type 2 diabetes pathogenesis. PLoS ONE 8, e55272 (2013).

31. A. Mahajan, et al., Fine-mapping type 2 diabetes loci to single-variant resolution using high-density imputation and islet-specific epigenome maps. Nat. Genet. 50, 1505–1513 (2018).

32. M. Vujkovic, et al., Discovery of 318 new risk loci for type 2 diabetes and related vascular outcomes among 1.4 million participants in a multi-ancestry meta-analysis. Nat. Genet. 52, 680–691 (2020).

33. A. Viñuela, et al., Genetic variant effects on gene expression in human pancreatic islets and their implications for T2D. Nat. Commun. 11, 4912 (2020).

34. L. Alonso, et al., TIGER: The gene expression regulatory variation landscape of human pancreatic islets. Cell Rep. 37, 109807 (2021).

35. J. Fadista, et al., Global genomic and transcriptomic analysis of human pancreatic islets reveals novel genes influencing glucose metabolism. Proc Natl Acad Sci USA 111, 13924–13929 (2014).

36. M. van de Bunt, et al., Transcript Expression Data from Human Islets Links Regulatory Signals from Genome-Wide Association Studies for Type 2 Diabetes and Glycemic Traits to Their Downstream Effectors. PLoS Genet. 11, e1005694 (2015).

37. A. Varshney, et al., Genetic regulatory signatures underlying islet gene expression and type 2 diabetes. Proc Natl Acad Sci USA 114, 2301–2306 (2017).

38. A. Karagiannopoulos, et al., Human pancreatic islet miRNA-mRNA networks of altered miRNAs due to glycemic status. iScience 25, 103995 (2022).

39. E. Que, et al., Genetic architecture modulates diet-induced hepatic mRNA and miRNA expression profiles in Diversity Outbred mice. Genetics 218 (2021).

40. H. Rutledge, et al., Identification of microRNAs associated with allergic airway disease using a genetically diverse mouse population. BMC Genomics 16, 633 (2015).

41. GTEx Consortium, et al., Genetic effects on gene expression across human tissues. Nature 550, 204–213 (2017).

42. P. Sethupathy, Illuminating microRNA Transcription from the Epigenome. Curr. Genomics 14, 68–77 (2013).

43. N. Sinnott-Armstrong, et al., Genetics of 35 blood and urine biomarkers in the UK Biobank. Nat. Genet. 53, 185–194 (2021).

44. Pan-UKB team, Pan-UK Biobank. https://pan.ukbb.broadinstitute.org (2020) (March 18, 2021).

45. T. Huan, et al., Genome-wide identification of microRNA expression quantitative trait loci. Nat. Commun. 6, 6601 (2015).

46. K. Sonehara, et al., Genetic architecture of microRNA expression and its link to complex diseases in the Japanese population. Hum. Mol. Genet. (2021) https://doi.org/10.1093/hmg/ddab361.

47. D. L. Taylor, et al., Integrative analysis of gene expression, DNA methylation, physiological traits, and genetic variation in human skeletal muscle. Proc Natl Acad Sci USA 116, 10883–10888 (2019).

48. O. Bruning, et al., Confounding Factors in the Transcriptome Analysis of an In-Vivo Exposure Experiment. PLoS ONE 11, e0145252 (2016).

49. O. Cabrera, et al., The unique cytoarchitecture of human pancreatic islets has implications for islet cell function. Proc Natl Acad Sci USA 103, 2334–2339 (2006).

50. M. Brissova, et al., Assessment of human pancreatic islet architecture and composition by laser scanning confocal microscopy. J. Histochem. Cytochem. 53, 1087–1097 (2005).

51. C. Ionescu-Tirgoviste, et al., A 3D map of the islet routes throughout the healthy human pancreas. Sci. Rep. 5, 14634 (2015).

52. G. Singh, K. B. Storey, MicroRNA Cues from Nature: A Roadmap to Decipher and Combat Challenges in Human Health and Disease? Cells 10 (2021).

53. M. A. Saunders, H. Liang, W.-H. Li, Human polymorphism at microRNAs and microRNA target sites. Proc Natl Acad Sci USA 104, 3300–3305 (2007).

54. M. E. Lacy, et al., Association of sickle cell trait with hemoglobin a1c in african americans. JAMA 317, 507–515 (2017).

55. G. Paré, et al., Novel association of HK1 with glycated hemoglobin in a non-diabetic population: a genome-wide evaluation of 14,618 participants in the Women’s Genome Health Study. PLoS Genet. 4, e1000312 (2008).

56. N. Soranzo, et al., Common variants at 10 genomic loci influence hemoglobin AL(C) levels via glycemic and nonglycemic pathways. Diabetes 59, 3229–3239 (2010).

57. P. Chen, et al., Multiple nonglycemic genomic loci are newly associated with blood level of glycated hemoglobin in East Asians. Diabetes 63, 2551–2562 (2014).

58. P. Chen, et al., A study assessing the association of glycated hemoglobin A1C (HbA1C) associated variants with HbA1C, chronic kidney disease and diabetic retinopathy in populations of Asian ancestry. PLoS ONE 8, e79767 (2013).

59. E. Wheeler, et al., Impact of common genetic determinants of Hemoglobin A1c on type 2 diabetes risk and diagnosis in ancestrally diverse populations: A transethnic genome-wide meta-analysis. PLoS Med. 14, e1002383 (2017).

60. K. L. Howe, et al., Ensembl 2021. Nucleic Acids Res. 49, D884–D891 (2021).

61. S. Fairley, E. Lowy-Gallego, E. Perry, P. Flicek, The International Genome Sample Resource (IGSR) collection of open human genomic variation resources. Nucleic Acids Res. 48, D941–D947 (2020).

62. A. Jones, et al., miRNA Signatures of Insulin Resistance in Obesity. Obesity (Silver Spring) 25, 1734–1744 (2017).

63. P. Wang, et al., miR-216a-targeting theranostic nanoparticles promote proliferation of insulin-secreting cells in type 1 diabetes animal model. Sci. Rep. 10, 5302 (2020).

64. D. Setyowati Karolina, S. Sepramaniam, H. Z. Tan, A. Armugam, K. Jeyaseelan, miR-25 and miR-92a regulate insulin I biosynthesis in rats. RNA Biol. 10, 1365–1378 (2013).

65. J. W. Kim, et al., miRNA-30a-5p-mediated silencing of Beta2/NeuroD expression is an important initial event of glucotoxicity-induced beta cell dysfunction in rodent models. Diabetologia 56, 847–855 (2013).

66. J. C. Kwekel, et al., Sex and age differences in the expression of liver microRNAs during the life span of F344 rats. Biol. Sex Differ. 8, 6 (2017).

67. M. N. Ziats, O. M. Rennert, Identification of differentially expressed microRNAs across the developing human brain. Mol. Psychiatry 19, 848–852 (2014).

68. S. Ameling, et al., Associations of circulating plasma microRNAs with age, body mass index and sex in a population-based study. BMC Med. Genomics 8, 61 (2015).

69. D. Lizarraga, et al., miRNAs differentially expressed by next-generation sequencing in cord blood buffy coat samples of boys and girls. Epigenomics 8, 1619–1635 (2016).

70. N. Mononen, et al., Whole blood microRNA levels associate with glycemic status and correlate with target mRNAs in pathways important to type 2 diabetes. Sci. Rep. 9, 8887 (2019).

71. J. López-Beas, et al., miR-7 Modulates hESC Differentiation into Insulin-Producing Beta-like Cells and Contributes to Cell Maturation. Mol. Ther. Nucleic Acids 12, 463–477 (2018).

72. J. M. Locke, G. da Silva Xavier, H. R. Dawe, G. A. Rutter, L. W. Harries, Increased expression of miR-187 in human islets from individuals with type 2 diabetes is associated with reduced glucose-stimulated insulin secretion. Diabetologia 57, 122–128 (2014).

73. R. Sato-Kunisada, N. Yoshida, S. Nakamura, H. Uchiyama, H. Matsumoto, Enhanced Expression of miR-199b-5p Promotes Proliferation of Pancreatic β-Cells by Down-Regulation of MLK3. Microrna 5, 57–65 (2016).

74. A. J. Lakhter, et al., Beta cell extracellular vesicle miR-21-5p cargo is increased in response to inflammatory cytokines and serves as a biomarker of type 1 diabetes. Diabetologia 61, 1124–1134 (2018).

75. L. Ding, et al., Identification of the differential expression of serum microRNA in type 2 diabetes. Biosci. Biotechnol. Biochem. 80, 461–465 (2016).

76. I. J. Gallagher, et al., Integration of microRNA changes in vivo identifies novel molecular features of muscle insulin resistance in type 2 diabetes. Genome Med. 2, 9 (2010).

77. S. Erener, et al., Deletion of pancreas-specific miR-216a reduces beta-cell mass and inhibits pancreatic cancer progression in mice. Cell Rep. Med. 2, 100434 (2021).

78. J. P. Werneck-de-Castro, M. Blandino-Rosano, D. Hilfiker-Kleiner, E. Bernal-Mizrachi, Glucose stimulates microRNA-199 expression in murine pancreatic β-cells. Journal of Biological Chemistry 295, 1261–1270 (2020).

79. S. Ibrahim, et al., β-Cell pre-mir-21 induces dysfunction and loss of cellular identity by targeting transforming growth factor beta 2 (Tgfb2) and Smad family member 2 (Smad2) mRNAs. Mol. Metab. 53, 101289 (2021).

