## Supplementary Appendix for "Human pancreatic islet microRNAs implicated in diabetes and related traits by large-scale genetic analysis"

### Supplementary Materials and Methods

#### Human pancreatic islet procurement and processing

We procured the human pancreatic islet (HPI) samples used in this study from the Integrated Islet Distribution Program, the National Disease Research Interchange (NDRI), or ProdoLabs. Tables S1,2 contain demographic and other reported information for the 69 samples (HPIs; age = 46.17 ± 10.71 y; 53.62% male; BMI = 27.49 ± 4.97 kg/$m^2$) considered in this study after quality control steps (see below). Islets were shipped overnight from the distribution centers. On receipt, we pre-warmed islets to 37°C in shipping media for 1-2 hours before harvest and harvested ~2,000-5,000 islet equivalents (IEQs) from each organ donor for subsequent molecular assays (e.g., RNA isolation).

#### Genotyping and quality control

We transferred 500-1,000 IEQs to tissue culture-treated flasks, cultured them as in Gershengorn et al. (1), and isolated genomic DNA for genotyping. We genotyped isolated DNA at the National Human Genome Research Institute Home (NHGRI) Genomics Core facility using either the HumanOmni2.5-4v1_H BeadChip array or the InfiniumOmni2-5Exome-8v1-3 BeadChip array (v1.3, Illumina, San Diego, CA, USA). Overall genotyping call rates ranged from 96.08-99.81%. As described previously (2), we mapped the Illumina array probe sequences to the GRCh37 (hg19) genome assembly using novoalign v2.07.11 (http://www.novocraft.com/products/novoalign). For downstream analysis and genotype imputation, we excluded SNPs with (i) ambiguous probe alignments, (ii) a 1000 Genomes (1000G) phase 3 variant with a minor allele frequency (MAF)≥1% within 7 base pairs (bp) of the 3' end of the probe, or (iii) call rates <95%—resulting in chip genotypes for 1,825,450 SNPs. Using SNP genotypes and KING v1.4 (3), we tested for sample relatedness and did not identify any individuals related at a 3rd-degree relationship or closer.

#### Sample ancestry and genotype PCs

To assess ancestry, we estimated genetic principal components (PCs) using weights produced with the Population Reference Sample (POPRES) reference panel (4) implemented in LASER (5). Briefly, we pruned genotyped, autosomal SNPs using plink v1.9 with a pairwise $r^2$ threshold of 0.5, excluding SNPs with a MAF≤1%, Hardy-Weinberg $P<10^{-6}$ (with options --indep-pairwise 50 5 0.5 --maf 0.01 --hwe 0.000001), or in regions of high linkage disequilibrium (LD) (6, 7). Next, we passed the genotypes of the remaining 564,920 SNPs to LASER to calculate genetic PCs of the samples in this study combined with the POPRES reference populations. We assigned ancestry based on the POPRES-predicted nearest neighbors (k=10). We identified 42, 7, and 6 individuals of European, African, and Hispanic/Latino ancestry, respectively.

To control for population stratification in downstream analyses, we performed a second round of principal component analysis (PCA) within each major ancestry present in this study (i.e., European, African, and Hispanic/Latino). For each major ancestry, we re-filtered SNPs following the method described above and used smartpca implemented in Eigensoft v6.1.4 (8, 9) to perform PCA with the retained SNPs. We used smartpca to perform the Tracy-Widom test (10) across the first 20 PCs and included PCs with $P\leq0.05$ in subsequent analyses. For European-only analyses, we used 460,072 filtered SNPs and identified 4 PCs. For African-only analyses, we used 208,609 filtered SNPs and identified 0 PCs. For Hispanic/Latino-only analyses, we used 146,562 filtered SNPs and identified 0 PCs.

#### Genotype imputation

We imputed genotypes as described in Lawlor et al. (2). Briefly, we assessed the allele frequency of each SNP by combining genotype data from the samples in this study with 140 samples from separate studies (2, 11) that were genotyped using similar array technology. We removed SNPs that had an alternate allele frequency difference >20% with the 1000G phase 3 EUR samples or that were palindromic with a MAF>20%, genotype missingness >2.5%, and Hardy-Weinberg $P<10^{-4}$. With the remaining 1,825,450 SNPs, we performed pre-phasing and imputation on autosomal markers using the Michigan Imputation Server (12). For pre-phasing, we used Eagle v2.3 (13) for autosomal chip markers and SHAPEIT v2.r790 (14) for chrX markers. We subsequently used minimac3 (12) for imputation of missing genotypes using the Haplotype Reference Consortium (hrc.r1.1.2016) panel (15). Finally, we removed all SNPs with an imputation $r^2 \leq 0.3$.

#### miRNA isolation, sequencing, and processing

We performed small RNA sequencing (smRNA-seq) to measure miRNAs using two different protocols: library preparation 1 (LP1) and library preparation 2 (LP2).

For LP1, we performed smRNA-seq as described in Lawlor et al. (2). Briefly, we extracted and purified total RNA from islets using Trizol (Life Technologies, Carlsbad, CA), generated smRNA libraries from 1 µg total RNA using Illumina's TruSeq Small RNA Library Kit, and performed single-end 51 base sequencing using an Illumina HiSeq2500 in 'Rapid Mode' with version 2 chemistry (Illumina, San Diego, CA).

For LP2, we isolated total RNA using the Total Purification kit (Norgen Biotek, Thorold, ON, Canada), following the manufacturer's instructions. We quantified RNA with a Nanodrop 2000 (Thermo Fisher Scientific, Waltham, MA) and determined RNA integrity using either an Agilent 2100 Bioanalyzer or a 4200 Tapestation (Agilent Technologies, Santa Clara, CA). We generated smRNA libraries using the CleanTag Small RNA Library Prep kit (TriLink Biotechnologies, San Diego, CA) and performed single-end 57 base sequencing using an Illumina HiSeq2500 (Illumina, San Diego, CA). Sequencing was performed at the Genome Sequencing Facility of the Greehey Children's Cancer Research Institute (University of Texas Health Science Center, San Antonio, TX).

For both preparations, we used miRquant v2.0 (16) to trim, map, and quantify smRNA-seq reads to the GRCh37 (hg19) genome assembly. We trimmed reads using Cutadapt v1.12 (17). Following miRquant guidelines, we discarded reads with <10 nucleotide overlap between the adapter sequence and the 3'-end of the sequencing read, ≥10% errors in the alignment, or <14 nucleotides in length after trimming. We aligned reads using Bowtie v1.1.0 (18) and created genomic windows containing miRNA loci from perfectly aligning reads. We re-aligned imperfectly mapped reads to these genomic windows to identify internal edits and non-templated nucleotide additions to the miRNAs using SHRiMP v2.2.2 (19). If a read aligned equally well to multiple genomic loci, we proportionally assigned reads to each of the loci. Finally, we quantified miRNA isoforms both as read counts (for differential gene expression analysis) and as normalized reads per million mapped to miRNAs (RPMMM; for genetic analyses). Given the uncertainty surrounding miRNA isoforms (isomiRs) shifted >2 nucleotides, we considered miRNAs with nucleotide shifts ≤2 in either direction as high confidence isomiRs for subsequent analyses.

#### miRNA quality control

For the analyses described in ***cis-miRNA-eQTL analysis***, ***Heritability estimates***, and ***Variance decomposition analysis*** and the PGS miRNA differential expression analysis described in ***miRNA differential expression analysis***, we were limited to the 64 samples from LP1 with both genotype and miRNA expression data. Using verifybamID v.1.1.1 (20), we compared miRNA reads aligned with exceRpt v4.4.0 (21) to the SNP chip genotypes and dropped 3 samples due to likely sample swaps (CHIPMIX>35%). We also dropped 4 samples due to low sequence coverage (<3 million reads). In total, we excluded 7 samples and used the remaining 57 samples for downstream analyses. For the for the analyses described in ***cis-miRNA-eQTL analysis***, ***Heritability estimates***, and ***Variance decomposition analysis***, we removed lowly expressed miRNAs by only keeping transcripts with raw counts≥100 in at least 25% of samples and kept only autosomal miRNAs, retaining 697 miRNAs. For the PGS miRNA differential expression analysis described in ***miRNA differential expression analysis***, we removed lowly expressed miRNAs by only keeping miRNAs with raw counts≥100 in at least 25% of samples in each major ancestry group, retaining 766, 727, and 661 miRNAs for European, African, and Hispanic/Latino ancestries, respectively.

For the T2D status, sex, age, and BMI miRNA differential expression analysis described in ***miRNA differential expression analysis***, we were not limited to samples with both genotype and miRNA expression data, so we used the 57 samples from LP1 and an additional 6 samples from LP2. Similar to the previous analyses, we removed lowly expressed miRNAs by only keeping transcripts with raw counts≥100 in at least 25% of samples, retaining 751 and 869 miRNAs from LP1 and LP2, respectively.

##### RNA isolation, sequencing, and processing

We performed RNA-seq as described in Lawlor et al. (2). Briefly, we extracted and purified total RNA from islets using Trizol (Life Technologies, Carlsbad, CA) and sequenced purified RNA using an Illumina NextSeq 500 with 2x101 bp cycles (Illumina, San Diego, CA). We aligned the processed reads to the GRCh37 (hg19) genome assembly using STAR v2.73a (22) with default parameters and quantified expression levels of Gencode v19 genes (Ensembl release 101) using QoRTs v1.3.6 (23). Finally, we quantified normalized mRNA expression as transcripts per million (TPM).

##### RNA-seq quality control

For the analyses described in ***Heritability estimates*** and ***Variance decomposition analysis***, we used 42 samples with both genotype and mRNA expression data. We used verifybamID v1.1.1 (20) to assess contamination of the total RNA used for RNA-seq in reference to the array genotypes in the coding regions of the genome. We excluded 1 sample that was likely contaminated (FREEMIX>15%) and 1 sample whose RNA did not match the corresponding genotypes (CHIPMIX>2%), resulting in 40 samples used for downstream analyses. We removed lowly expressed mRNA by only keeping genes with raw counts≥100 in at least 25% of samples, retaining 14,805 autosomal mRNAs.

##### *cis*-miRNA-eQTL analysis

We performed a *cis*-miRNA-eQTL analysis by testing all SNPs within 250 kilobases (kb) of the mature miRNA transcript using LIMIX v3.0.4 (24) and the 57 samples with imputed genotypes and miRNA. For genotypes, we selected 4,741,068 bi-allelic, autosomal SNPs with minor allele count (MAC)>10 across the 57 samples. For miRNAs, we removed miRNAs that were lowly expressed across the 57 samples as described in ***miRNA quality control***, retaining 697 miRNAs for analysis.

To account for unknown covariates affecting miRNA expression, we used PEER v1.3 (25) to infer and adjust the miRNA expression data for 5 hidden factors. For PEER correction, we included age and sex as known covariates. As performed by previous studies (26, 27), we inverse rank-normalized the residual values from PEER, where the effects of known and discovered factors had been removed.

To account for effects of population structure or cryptic relatedness, we calculated the genetic relatedness matrix (GRM) across all 4,741,068 SNPs as described in Hoffman (28) and included the GRM as a random effect term, *K*, in subsequent models. Briefly, let $X_{ij}$ represent the scaled, centered genetic dosage for individual *i* at SNP *j*. Thus, $K=XX^T$. For the eQTL analysis, we calculated the normalized GRM, $K^* = K / mean(diag(K))$.

For each miRNA, we considered the following generalized linear mixed models:

$$Y = M\alpha + K^{*}\upsilon + \varepsilon \qquad (1)$$

$$Y = M\alpha + G\beta + K^{*}\upsilon + \varepsilon \quad (2)$$

where $Y$ is the inverse rank-normalized PEER residuals described above, $M$ is a matrix of fixed-effect covariates (here, a vector of ones as an offset term), $G$ is a matrix of candidate genetic variants, $K^*$ is the normalized GRM as defined above, ε is a noise variable, and α, β, and υ are the corresponding regression coefficients. Notably, equation (1) is effectively equation (2), where we assume no genetic effects, β=0. To assess the association between genetic effects and the phenotype, we derived *P*-values by performing a likelihood ratio test using the marginal likelihoods of the two models (24).

To control for correlation among SNPs, we calculated empirical *P*-values ($P_{emp}$) per miRNA, similar to Ongen et al. (29). Briefly, for each miRNA, we randomly permuted the sample ids of $G$ and re-ran the *cis*-miRNA-eQTL analysis 10,000 times. Using these permuted *P*-values, we calculated $P_{emp}$ as the proportion of permuted *P*-values less than or equal to the minimum nominal *P*-values. Finally, we controlled the false discovery rate (FDR) across all tested miRNAs using Storey's method (30) and $P_{emp}$.

**Heritability estimates**

For miRNAs and mRNAs, we estimated the SNP-based heritability ($h_g^2$)—the proportion of variation in expression that can be explained by additive effects of commonly occurring genetic variants for each transcript—by fitting the following model using LIMIX v3.0.4 (24):

$$Y = M\alpha + K^{*}\nu + \varepsilon \quad (3)$$

where the terms are the same as defined in equations (1) and (2). We considered highly heritable transcripts to have $v \geq 0.9$.

For miRNAs, we used the paired genotype and miRNA data as defined in ***cis-miRNA-eQTL analysis***. For mRNAs, we used 40 samples with both imputed genotypes and mRNA expression data that passed the QC filters described in ***RNA-seq quality control***. We included 4,040,664 bi-allelic, autosomal SNPs with MAC>10 across the 40 samples to calculate $K^*$. For both the miRNA and mRNA datasets, we used the methodology described in ***cis-miRNA-eQTL analysis*** to account for unknown covariates in expression data, including age and sex as covariates. Briefly, we used PEER to adjust expression data for 5 hidden factors and calculated the inverse rank-normalized residual values of the adjusted expression data to use in heritability estimates.

**Variance decomposition analysis**

To investigate the contributions of *cis*- and *trans*-acting genetic effects to miRNA and mRNA expression, we performed variance decomposition using LIMIX v3.0.4 (24). Let $K^*$ denote the normalized GRM, $K$. For each transcript, we fit the following equation:

$$Y = M\alpha + K^{*}_{cis} v_{cis} + K^{*}_{trans} v_{trans} + \varepsilon \quad (4)$$

where $K_{cis}$ is the GRM calculated using SNPs within 250kb of either side of the mature miRNA body or mRNA transcription start site (TSS; presumed to be *cis*-acting), $K_{trans} = K_{all} - K_{cis}$, and the rest of the terms are the same as equations (1) and (2). To represent the proportion of variance explained by a single component, we used the following formula:

$$h_x = \frac{v_x}{v_x + v_{S-x} + \varepsilon} \quad (5)$$

where $S = \{ cis, trans \}$ and $x \subset S$.

For both miRNA and mRNA, we used the PEER-adjusted expression data described in ***Heritability estimates***. To account for the fact that some miRNA TSSs may be quite far away from the miRNA body, we also re-estimated genetic effects using a larger 20Mb window to define the set of *cis*-acting SNPs.

#### Genetic colocalization analysis

For each miRNA-eQTL, we performed colocalization analysis using coloc v3.1 (31) with default priors and all variants present in both the miRNA-eQTL analysis and summary statistics of the genetic analysis of interest. Briefly, coloc outputs posterior probabilities of association for five different hypotheses: PP0 (neither trait is associated), PP1 (only the miRNA-eQTL is associated), PP2 (only the alternative trait is associated), PP3 (both traits are associated with different causal variants), and PP4 (both traits are associated with the same causal variant). Based on Guo et al. (32), we considered two genetic signals to have strong evidence of colocalization if PP3+PP4≥0.99 and PP4/PP3≥5 and weak evidence of colocalization if PP3+PP4≥0.9 and PP4/PP3≥3. For disease and quantitative traits, we used summary statistics from common variant genetic association studies for fasting blood glucose levels (33), blood glucose levels (34), fasting insulin (33), glycated hemoglobin (HbA1c) (34), T2D (35), triglycerides (TG) (34), and red blood cell distribution width (RCDW) (36). To identify potential downstream targets of miRNAs, we also colocalized each miRNA-eQTL with exon- and gene-level mRNA eQTL results in HPIs (37). Finally, we only considered miRNA-eQTLs that colocalized with genetic studies of T2D and glycemic traits, where at least one study had a lead SNP *P*-value$\leq 5\times10^{-8}$ at the miRNA-eQTL locus.

#### Mediation analysis

We sought to determine whether miRNA expression may causally influence mRNA expression for miRNA-mRNA colocalization pairs found in ***Genetic colocalization analysis*** using CIT v2.2 (38) as described in Taylor et al. (39). We used the 34 samples with paired genotype, miRNA expression, and mRNA expression data. For both miRNAs and mRNAs, we used the PEER-adjusted expression data described in ***Heritability estimates***.

Briefly, CIT performs a series of conditional regression tests to determine if a genetic effect (instrument) on a trait (outcome) is mediated by another molecular effect (exposure). For each miRNA-mRNA colocalization pair, we considered miRNA expression as the exposure, mRNA expression as the outcome, and the miRNA-eQTL tag SNP as the instrument. Thus, the CIT model corresponded to SNP→miRNA expression→mRNA expression. We performed multiple hypothesis correction using Bonferroni correction over the number of miRNA-mRNA colocalization pairs, *m*, and considered miRNA-mRNA colocalization pairs with $P\leq0.05/m$ to be evidence of miRNA-mediated mRNA eQTLs.

#### *trans*-eQTL analysis

We tested for associations between tag variants from the miRNA-eQTL results and mRNA expression using LIMIX v3.0.4 (24) as described in ***cis-miRNA-eQTL analysis***. Briefly, for each mRNA, we modeled the expression with and without the genetic effect of interest. In both models, we included an offset term as a fixed-effect covariate, the GRM as a random-effect covariate to account for hidden effects of population structure or cryptic relatedness, and a term for noise. Next, we performed the likelihood ratio test of the two models to derive nominal *P*-values of association. Finally, we controlled the FDR across all tested variant-gene pairs using Storey's method (30) and the nominal *P*-values.

We used the 40 sample dataset with paired mRNA expression data across 14,805 mRNAs and imputed genotypes as described in ***Heritability estimates***, including the filtered 4,040,664 bi-allelic, autosomal SNPs to calculate the GRM. To account for known and unknown covariates, we also used the same inverse rank-normalized residual values of the PEER-adjusted mRNA expression as the phenotype in the LIMIX model.

#### Comparison of HPI miRNA-eQTLs and blood miRNA-eQTLs

We compared SNP-miRNA effect sizes in HPI miRNA-eQTLs to miRNA-eQTL results from previously reported genetic analyses in blood (40, 41). For Huan et al., we downloaded the published summary statistics, which included all tested SNP-miRNA pairs at an FDR≤10%. For Sonehara et al., we downloaded the published summary statistics, which included all tested SNP-miRNA pairs with $P \leq 0.05$ (National Bioscience Database Center (NBDC) Human Database accession number hum0197.v6). For each comparison (i.e., this study vs. Huan et al. and this study vs. Sonehara et al.), we retained shared SNP-miRNA pairs and harmonized the effect alleles by reversing the effect size estimates of blood SNP-miRNA pairs if the effect alleles were swapped. We determined the correlation of the effect size estimates between studies by calculating Spearman's rank correlation coefficient.

#### Allelic imbalance in smRNA-seq

We tested for allelic imbalance in the smRNA-seq data at heterozygous SNP sites in the 57 islet samples with both smRNA-seq and imputed genotypes (see ***miRNA isolation, sequencing, and processing***) using the method described in Greenwald et al. (42). Briefly, we used the mem function from bwa v0.7.17-r1194-dirty (43) with options "-q 30 -F 2816" to re-align single-end smRNA-seq reads to the GRCh37 (hg19) reference genome and filter reads with low mapping quality. For each sample, we used WASP v0.3.4 (44) to remove duplicate reads and correct for reference mapping bias. Next, we tested for allelic imbalance in SNPs where we had at least two heterozygotes with two or more reads covering each allele by performing a binomial test in each sample. We used the resulting *P*-values to calculate signed Z-scores and combined the Z-scores across samples using Stouffer's Z-score method and sequencing depth to weigh each sample. Finally, we performed multiple hypothesis correction using the Benjamini-Hochberg procedure (45) and *P*-values derived from the combined Z-scores.

#### Intersection of T2D-related SNPs with mature miRNA bodies and target regions

To investigate the overlap of T2D-related SNPs and miRNA target regions within mRNA transcripts, we used the intersect function from bedtools v2.29.2 (46) to identify overlaps between the genomic coordinates of predicted miRNA target sites from TargetScan v7.2 (47) and genetic variants from 99% credible sets from genetic studies for T2D (35), HbA1c (34), and blood glucose (34). In some cases, the miRNA target site was split across two exons, so the genomic coordinates contained introns. We used GRCh37 (hg19) exon coordinates to remove SNPs lying in intronic regions (i.e., SNPs that would not affect miRNA binding to the 3' UTR site). To identify the most islet-relevant miRNA target sites, we only kept SNPs where the miRNA and target mRNA were highly expressed by filtering for transcripts with raw counts≥100 in at least 25% of samples. We used the 57 samples in LP1 (described in **miRNA quality control**) for the miRNA filter and the 40 samples with mRNA expression data (described in **RNA-seq quality control**) for the mRNA filter. Finally, to evaluate the effect of SNPs on target mRNA levels, we used conditional eQTL summary statistics from a study of HPIs from 420 individuals (37) for mRNAs identified with T2D-related SNPs in miRNA target regions. Taking the tag SNPs for these eQTLs, we calculated LD with the T2D-related SNPs in miRNA target regions in European populations (i.e., 1000GENOMES phase_3 FIN, CEU, GBR populations) (48, 49) and considered those with $r^2 > 0.9$.

To investigate the overlap of T2D-related SNPs and mature miRNAs, we performed the procedure outlined for target regions but using the mature miRNA genomic coordinates from miRbase (release 18) (50–55). Briefly, we used bedtools v2.29.2 (46) to intersect the mature miRNA genomic coordinates with the same T2D-related 99% credible sets and kept SNPs where the miRNA transcript was highly expressed (raw transcript counts≥100 in at least 25% of samples). For the miRNA filter, we used the 57 samples in LP1 (described in **miRNA quality control**). Since we identified no overlaps, we did not perform further analyses.

#### Modeling an interaction effect between rs1464569 and miR-532-3p expression on *NICN1* expression

To investigate the possibility of an interaction effect between rs1464569 and miR-532-3p expression on *NICN1* expression, we used the qtl.iscan function from LIMIX v3.0.4 (24) and the 34 samples with imputed genotypes, miRNA expression data, and mRNA expression data. We used the RPMMM-normalized counts for miR-532-3p expression and the TPM-normalized counts for *NICN1* expression. To account for effects of population structure or cryptic relatedness, we calculated the normalized GRM, $K^*$, as described in ***cis-miRNA-eQTL analysis*** using the 3,701,248 bi-allelic, autosomal SNPs with MAC>10 across the 34 samples.

We considered the following generalized linear mixed models:

$$Y = M\alpha + K^{*}\upsilon + \varepsilon \qquad (6)$$

$$Y = M\alpha + (G * E)\beta + K^{*}\upsilon + \varepsilon \qquad (7)$$

where $Y$ is the inverse rank-normalized *NICN1* expression, $G*E$ is the interaction between rs1464569 ($G$) and the inverse rank-normalized miR-532-3p expression ($E$), and the remaining terms are the same as defined in equations (1) and (2). As covariates, we passed sex (with male as reference), age, rs1464569 dosages, inverse rank-normalized miR-532-3p expression, and a vector of ones as an offset term. We performed a likelihood ratio test using the marginal likelihoods of the two models to derive the nominal *P*-value (24). To derive an empirical *P*-value, we performed the permutation analysis described in ***cis-miRNA-eQTL analysis***. Briefly, we randomly permuted the sample ids of *G* and re-ran the interaction analysis 10,000 times. Using these permuted *P*-values, we calculated $P_{emp}$ as the proportion of permuted *P*-values less than the nominal *P*-value.

To evaluate the effect of cell type composition, we repeated the above analysis, incorporating the cell type proportions calculated in ***Cell type composition estimation*** as an additional covariate. We found that incorporating cell type proportions had no effect on the results.

#### Polygenic score calculations

We calculated polygenic scores for each genotyped participant using the --score function from PLINK v1.9 (56, 57) and SNP weights derived from genetic associations for T2D (35), fasting blood glucose (33), fasting insulin (33), blood glucose (34), and HbA1c (34). For the SNP weights, we used previously reported weights for blood glucose (34, 58) and HbA1c (34, 58) and calculated weights using PRS-CS (release Sept. 10, 2020) (59) for the remaining phenotypes with European samples from the 1000 genomes project as a linkage disequilibrium reference panel.

#### miRNA differential expression analysis

We tested for association of miRNA expression with T2D status, sex, age, BMI, and PGSs for T2D and glycemic traits (described in ***Polygenic score calculations***). Depending on the phenotype, we fit different models to account for dataset specific effects. For T2D status, sex, age, and BMI, we analyzed miRNA data from three library preparation (LP) methods: LP1 from this study (n=2 T2D, n=55 NGT; 751 miRNAs), LP2 from this study (n=2 T2D, n=4 NGT; 869 miRNAs), and a third LP method from a publicly available miRNA expression dataset (n=4 T2D, n=3 NGT; 295 miRNAs) (60). To control for differences across library preparation methods, we fit separate models for each LP and meta-analyzed the results. For the Kameswaran et al. dataset, we used the same processing pipeline that we used for LP1 and LP2: we quantified miRNA counts from fastQ files using the miRquant pipeline (see ***microRNA isolation, sequencing, and processing***) and filtered lowly-expressed miRNAs by retaining miRNAs with raw counts≥100 in at least 25% of samples (see ***microRNA-seq quality control***). For the PGSs, we used the 57 samples from LP1 which had both genotype and miRNA data. To account for ancestry specific biases in PGSs (61), we fit separate models for each major ancestry—European (n=42; 766 miRNAs), African (n=7; 727 miRNAs), and Hispanic/Latino (n=6; 661 miRNAs)—and meta-analyzed the results.

For each phenotype, we used a three-stage pipeline to identify differentially expressed (DE) miRNAs within each dataset (i.e., library preparation or ancestry). We (i) performed a first-pass Wald test using DESeq2 v1.32.0 (62), (ii) identified factors of unknown variation using RUVSeq v1.26.0 (63) and empirically derived control genes from stage i, and (iii) re-ran stage i and incorporated the calculated RUVSeq factors from stage ii as additional covariates. To determine the final set of phenotype-associated miRNAs, we meta-analyzed the results across datasets using MetaVolcano v1.6.0 (64).

In stage i, we used DESeq2 to pre-compute size factors across samples with the pre-filtered miRNA expression counts to capture the full library sizes of each sample. Then, we tested for DE miRNAs using DESeq2's Wald test with default parameters, raw miRNA counts (described in ***miRNA quality control***), the pre-computed size factors, and a set of covariates specific for each phenotype and dataset (Table S5,6). Across all phenotypes, we included sex and age as covariates, where age was standardized to unit variance (mean centered and scaled). For the age, sex, and BMI tests, we included T2D status as an additional covariate. For BMI, we mean-imputed three samples with missing BMI values and standardized these values to unit variance (mean centered and scaled). For the PGS analyses, we also included the genetic PCs as additional covariates according to the analysis described in ***Sample ancestry and genotype PCs*.** Briefly, we included 4 PCs for the European ancestry analyses and 0 PCs for the other ancestries (African and Hispanic/Latino), since the PC analysis reported little variation in samples of African and Hispanic/Latino ancestry. For some models (e.g., BMI analysis in the Kameswaran et al. dataset), we were restricted by sample size in the number of terms that we could include in the model (the number of covariates included in a model cannot equal the total number of samples being considered). In such cases, we prioritized dropping the age covariate from the model as we identified no age-associated miRNAs.

In stage ii, we performed RUVSeq analysis, a method designed to control for unknown variation in differential expression studies. Briefly, in order to run RUVSeq, one must define a set of control transcripts that are not correlated with the phenotype of interest. Following the RUVSeq recommendations, we used the results from stage i to identify control miRNAs, defined as the half with the highest $P$, excluding miRNAs with $P \leq 0.25$. We used the DESeq2-normalized counts of the control miRNAs and the RUVg function from RUVSeq to calculate two factors of unwanted variation.

In stage iii, we repeated the stage i analysis and included the 2 RUVSeq factors from stage ii as additional covariates. For some models, we dropped a RUVSeq factor as a covariate due to the small sample size of the dataset and the limitations that sample size imposes on the number of covariates that one can model (Table S5,6).

Finally, for all miRNAs observed in at least 2 datasets, we meta-analyzed the effects across datasets using a random effect model implemented by the rem_mv function in MetaVolcanoR with default parameters. We attempted to remove miRNAs that showed effect size heterogeneity using Cochran's *Q*-test (65) as implemented in MetaVolcanoR, but we found no miRNAs with $P_{Cochran} \leq 0.05$. For miRNAs observed in a single dataset, we retained the original stage iii *P*-values. We performed multiple hypothesis correction using Storey's method (30) and considered miRNAs with $|\log_2(FC)| \geq 1$ and FDR≤0.05 to be differentially expressed.

##### Cell type composition estimation

Using fastQ files from a publicly available miRNA expression dataset of sorted human alpha and beta cells (60), we quantified miRNA counts using the miRquant pipeline described in ***miRNA isolation, sequencing, and processing***. To deconvolute cell type composition, we used 532 miRNAs that had raw counts≥100 in at least one of the cell types and were also present in the samples from this study with average raw counts≥100. We estimated the cell composition of islets in this study using the unmix function from DESeq2 v1.30.1 (62) (Fig. S16). We ran unmix with default parameters and defined the alpha value with DESeq2's dispersionFunction using fitType = "parametric". We note the limitation that there are islet cell types that are unaccounted for in this deconvolution (e.g., delta cells) as we did not have miRNA expression profiles for these cell types.

To evaluate the effect of cell type composition in the presented data, we repeated analyses and incorporated cell type composition as an additional covariate. For ***cis-miRNA-eQTL analysis***, we included cell type composition as a covariate during PEER correction. For ***miRNA differential expression analysis***, we included cell type composition as a covariate in the DESeq2 model. Due to limited sample size among specific library preparations and ancestries, we only compared the effect of cell type composition in the LP1-specific DE miRNAs for the sex, age, BMI, and T2D status analyses and European-specific DE miRNAs for the PGS analyses. We found that incorporating cell type composition had a minimal impact on the final results (Fig. S17).

##### Data availability

The individual-level genotype, RNA-seq, and LP1 smRNA-seq data generated in this study are available in the database of Genotypes and Phenotypes (dbGaP) accession number phs001188.v2.p1 and are accessible through dbGaP's standard data access request procedures. The LP2 smRNA-seq are available in the Gene Expression Omnibus (GEO) data repository under the GSE196797 accession number.

#### Supplementary Figures

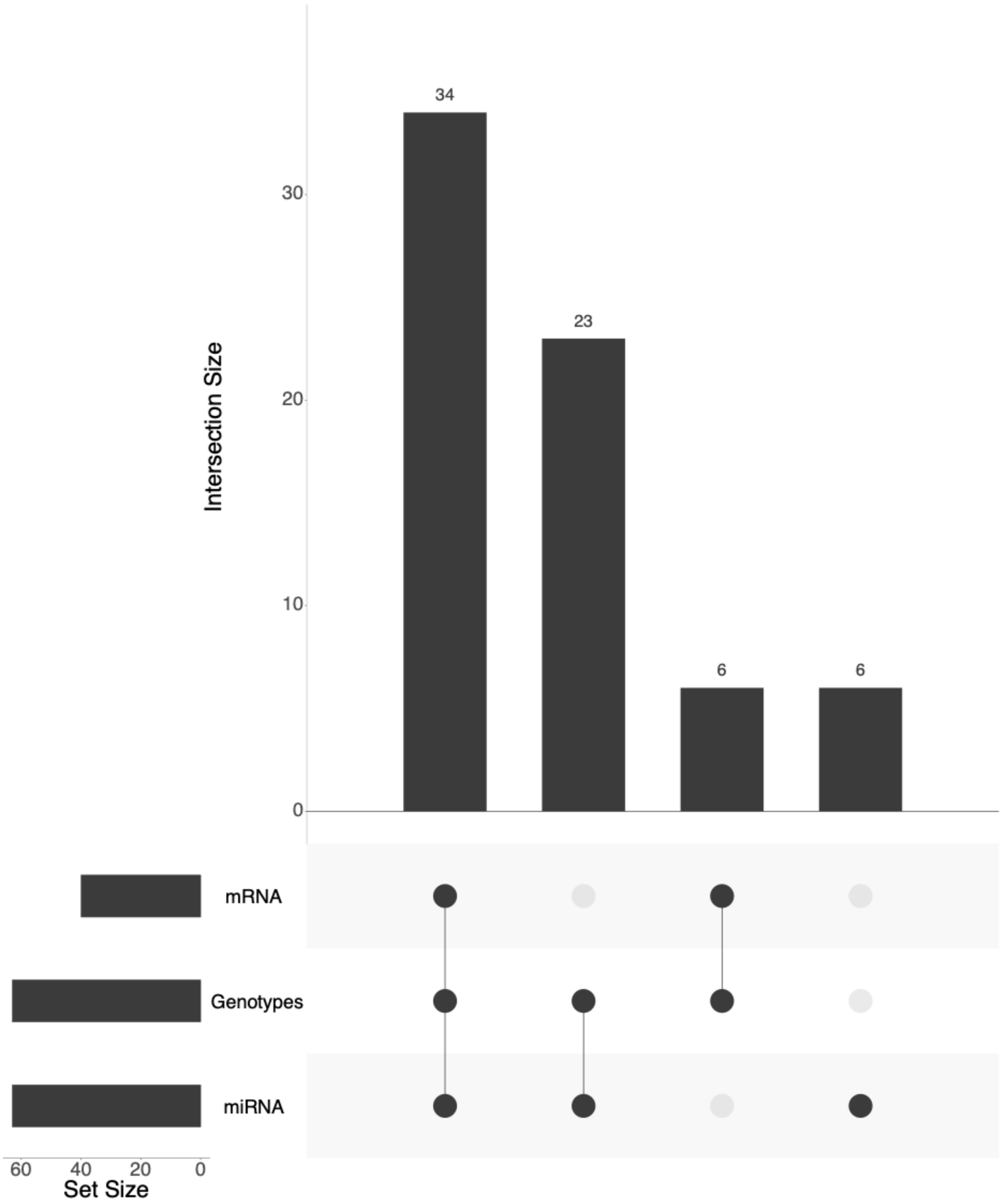


**Fig. S1. Data modality overview.** Number of samples with data from each modality.

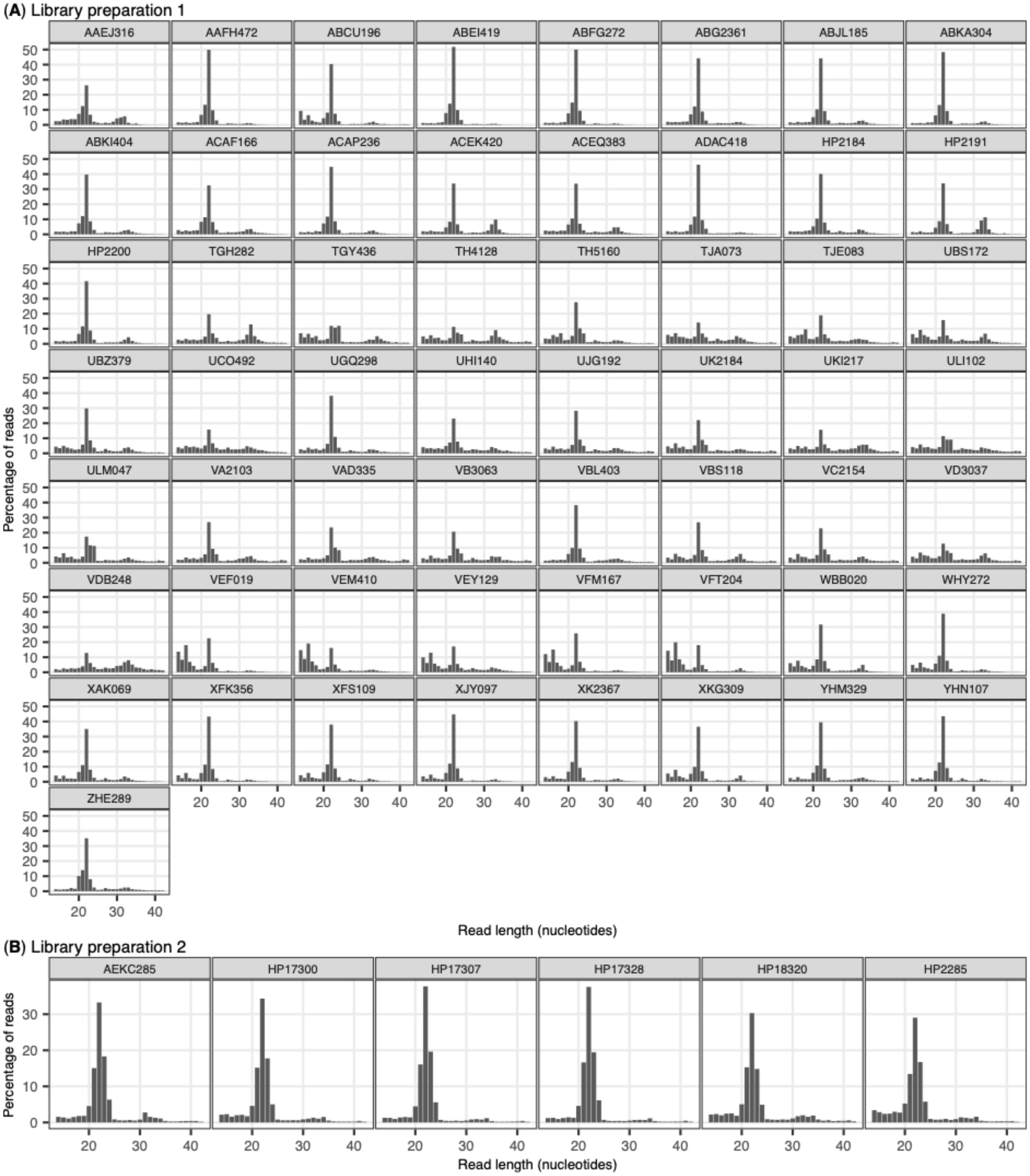


**Fig. S2. Read length distribution of smRNA-seq reads.** Read length distribution of smRNA-seq reads across samples.

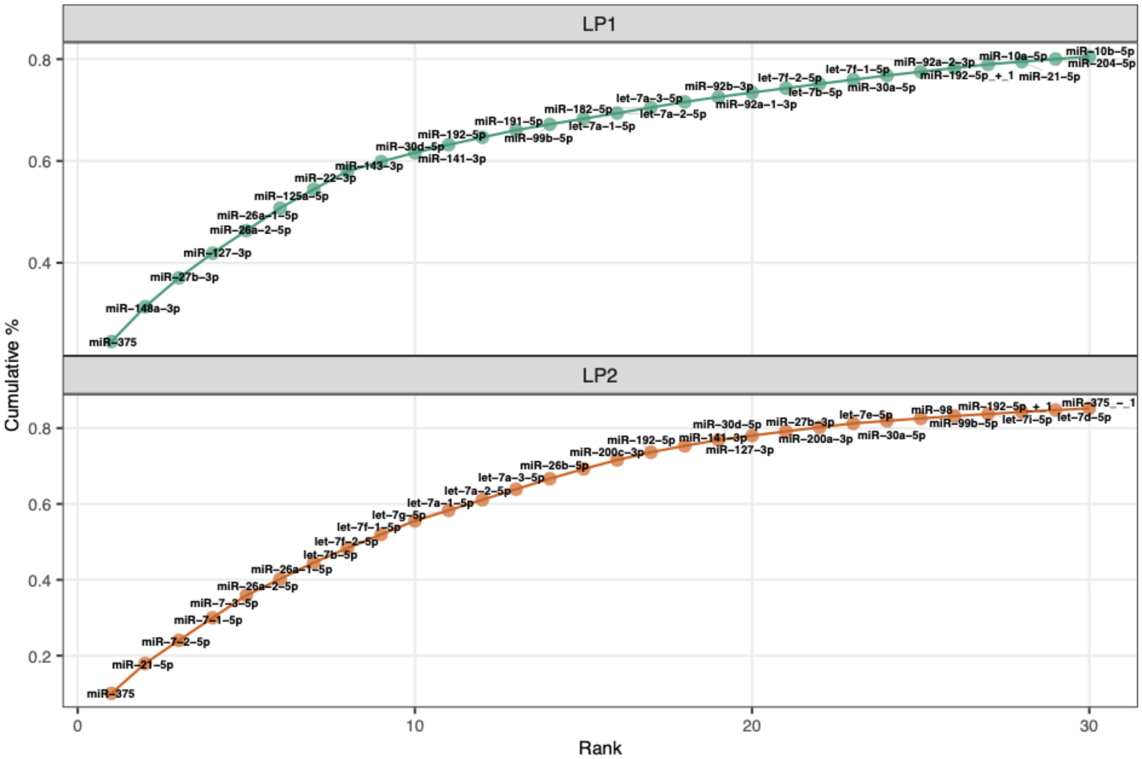


**Fig. S3. Cumulative miRNA counts across library preparations.** For each library preparation (i.e., library preparation 1 (LP1) and library preparation 2 (LP2)), miRNA counts are summed across samples.

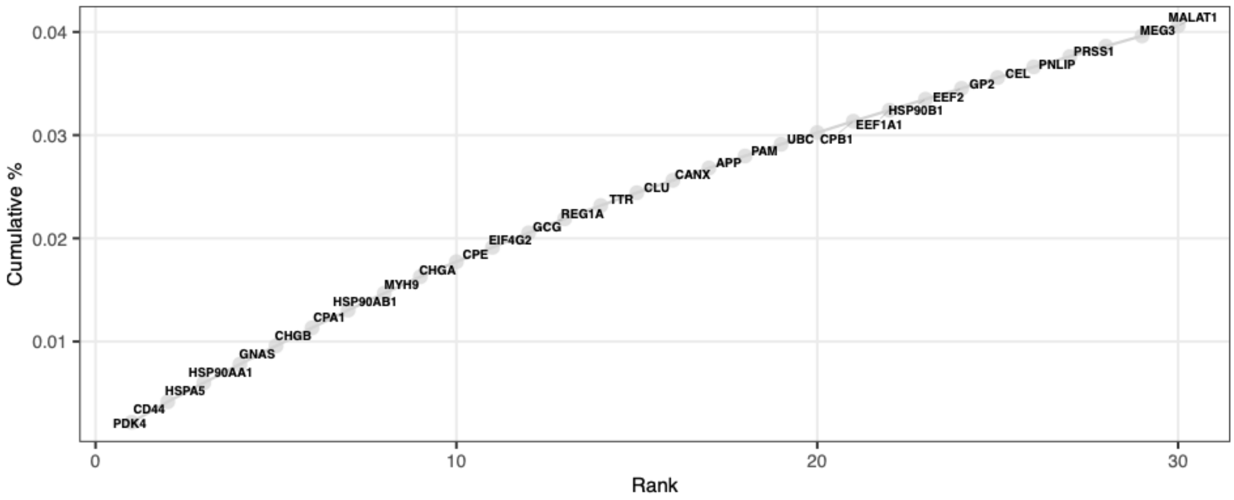


**Fig. S4. Cumulative mRNA counts.** mRNA counts are summed across samples.

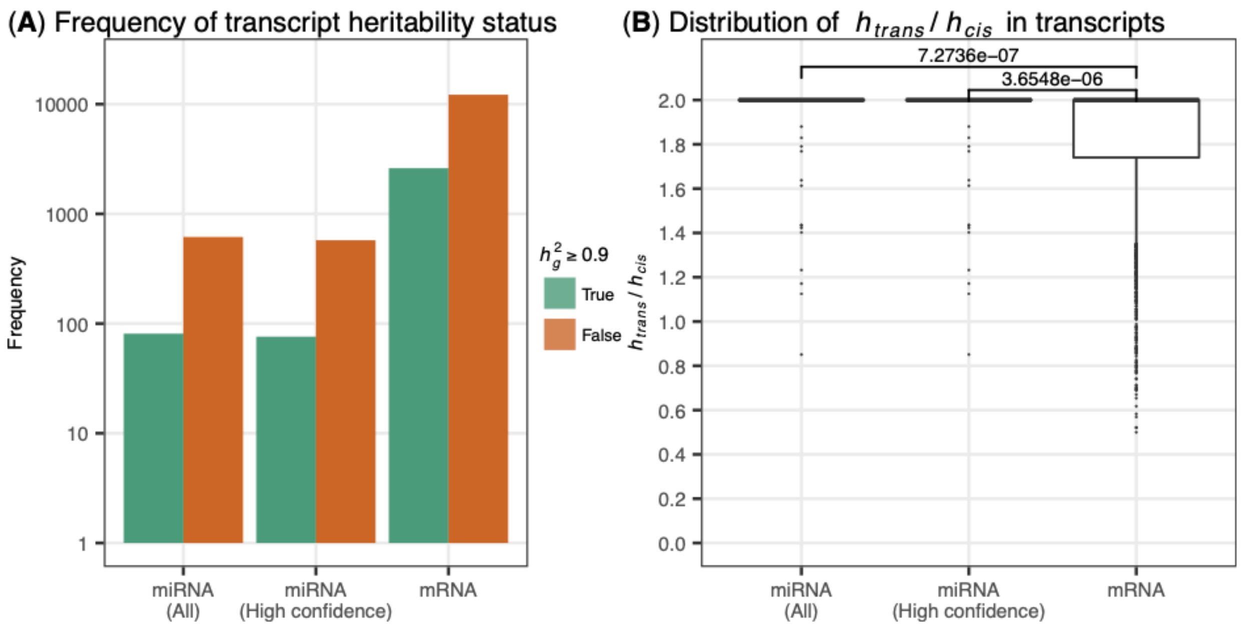


**Fig. S5. Heritability and variance decomposition results in miRNA and mRNA expression data.** (A) Number of heritable transcripts ($h_g^2 \geq 0.9$) across miRNAs and mRNAs. (B) Distribution of the ratio between variance explained by *trans*-effects ($h_{trans}$) to variance explained by *cis*-effects ($h_{cis}$) in heritable transcripts. *P*-value calculated using Mann-Whitney rank test.

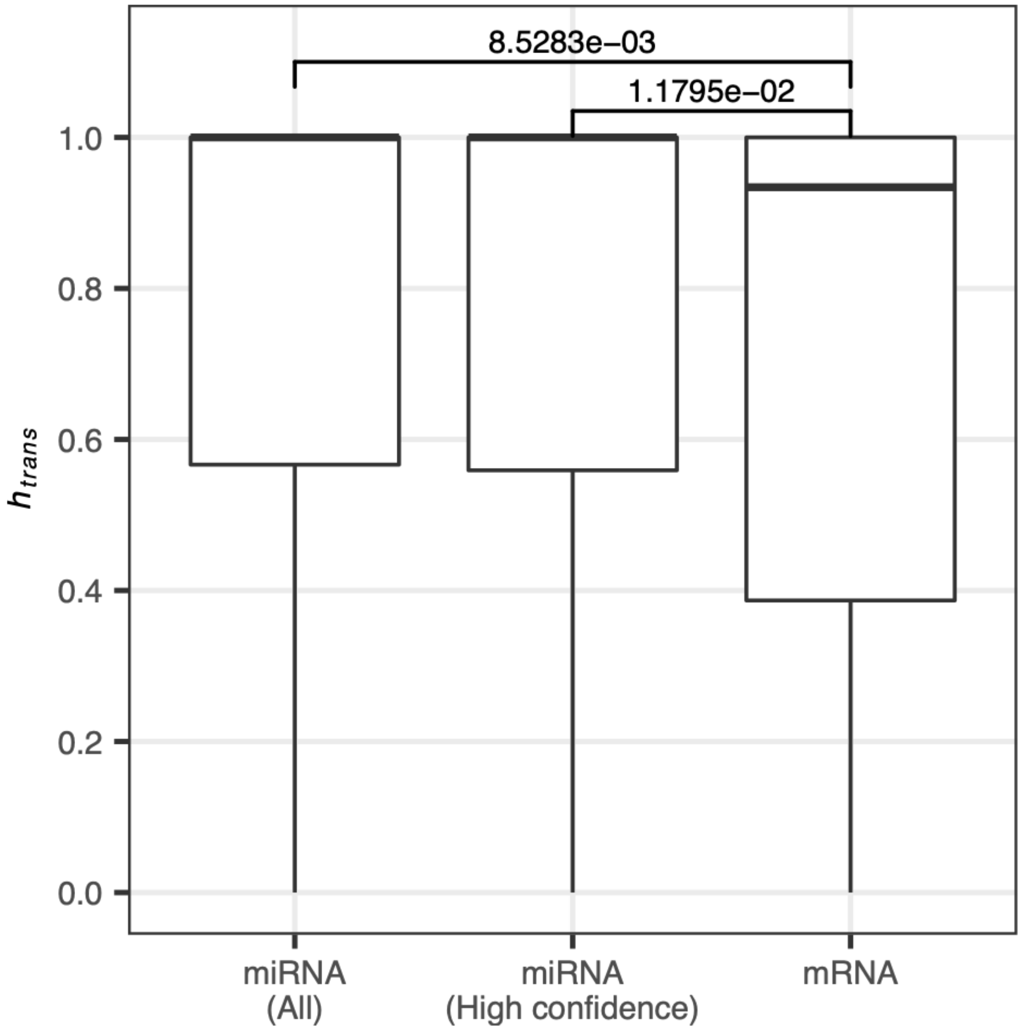


**Fig. S6. Variance decomposition results in miRNA and mRNA expression data using a 20Mb window to define *cis*-effects.** Distribution of variance explained by *trans*-acting genetic effects in expression of heritable transcripts. *P*-value calculated using Mann-Whitney rank test.

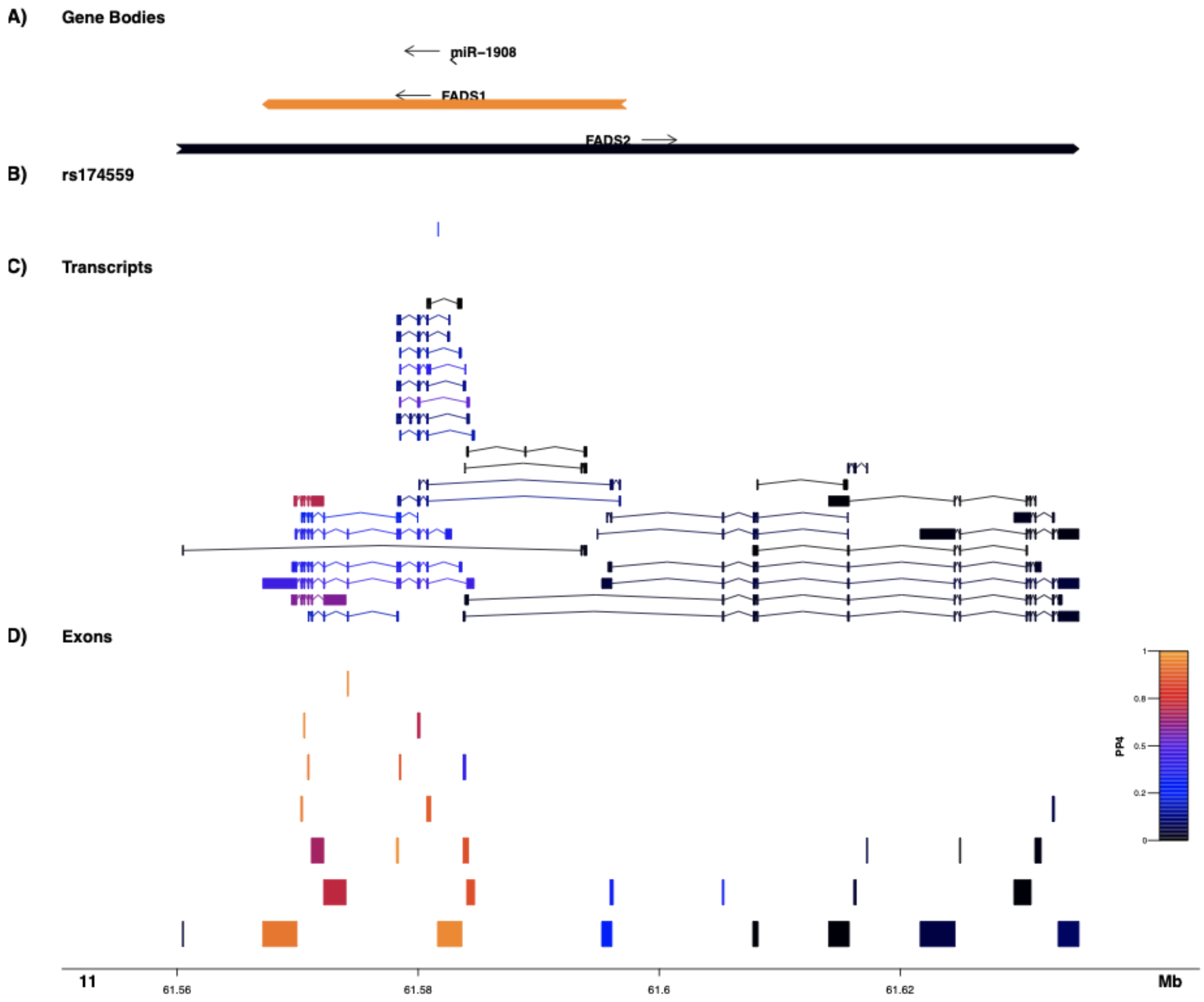


**Fig. S7. miR-1908 eQTL colocalization results with exon- and gene-level eQTLs from human pancreatic islets.** Evidence of genetic colocalization between miR-1908 eQTL and genetic association signals for exon and gene expression in human pancreatic islets (see Methods). Features are colored by PP4 (i.e., posterior probability that both traits are associated and share a single causal variant). (A) Gene bodies features. (B) Position of the rs174559 tag SNP for the miR-1908 eQTL. (C) Transcripts of gene bodies. (D) Exons of gene bodies.

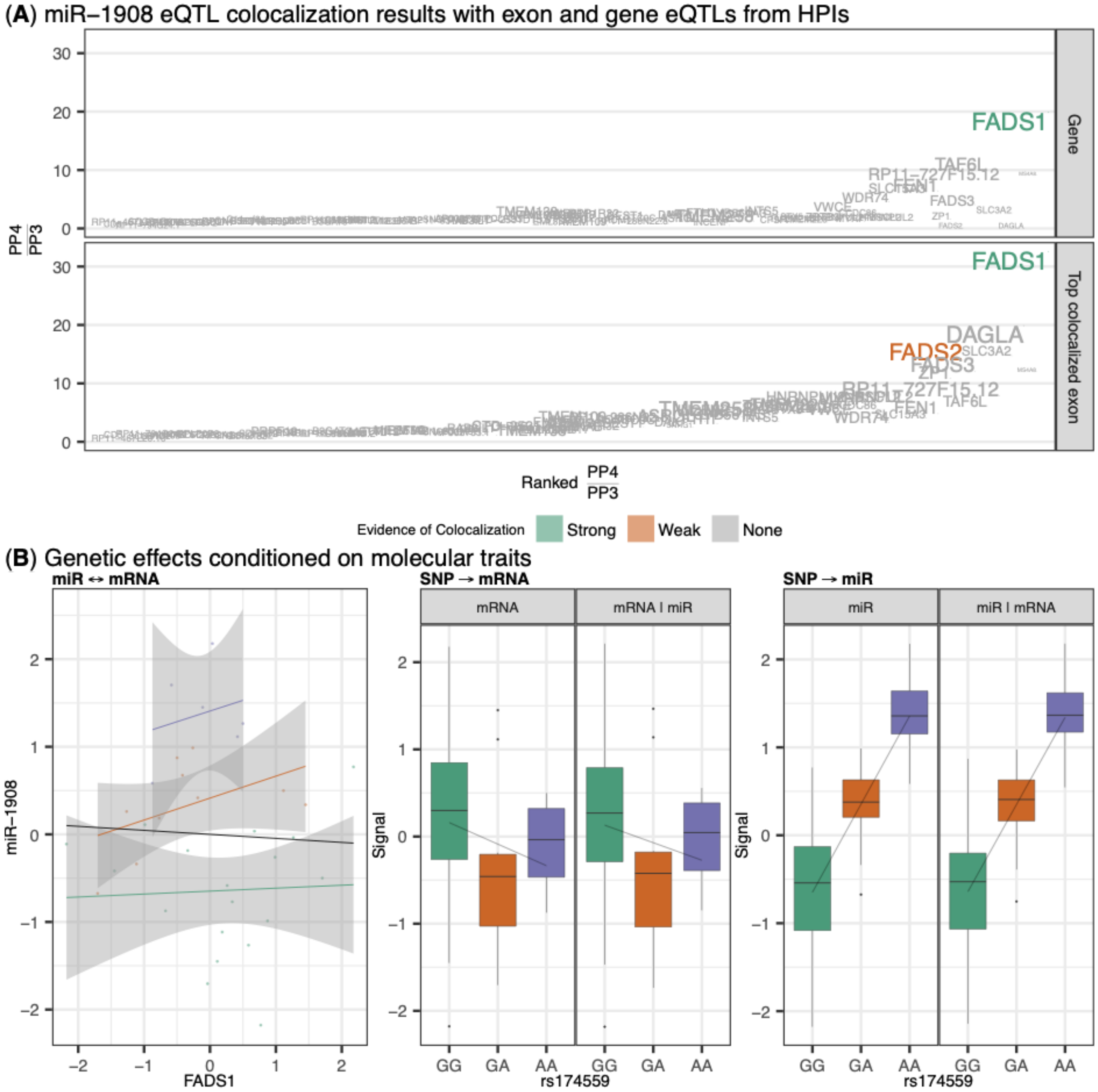


**Fig. S8. Identifying downstream targets of miR-1908.** (A) Evidence of genetic colocalization between miR-1908 eQTL and genetic association signals for exon and gene expression in human pancreatic islets (HPIs; see Methods). Posterior probability definitions: PP3 - both traits are associated with different causal variants, PP4 - both traits are associated and share a single causal variant. (B) Results from mediation analysis between miR-1908, *FADS1*, and rs174559 (see Methods). Left: scatter plot of residual *FADS1* expression (adjusted for PEER factors used in $h_g^2$ analysis; x-axis) and residual miR-1908 expression (adjusted for PEER factors used in miRNA-eQTL analysis; y-axis). Linear regression line for gene expression overall (black) and colored by the rs174559 genotype (GG, green; GA orange; AA, purple). Middle: box plots and linear regression line (additive model) of residual *FADS1* expression by rs174559 genotype (mRNA facet). Box plot and regression line of residual *FADS1* expression by rs174559 genotype after adjustment for residual miR-1908 expression (mRNA|miR facet). Right: box plots and linear regression line (additive model) of residual miR-1908 expression by rs174559 genotype (miR facet). Box plot and regression line of residual miR-1908 expression by rs174559 genotype after adjustment for residual *FADS1* expression (miR|mRNA facet).

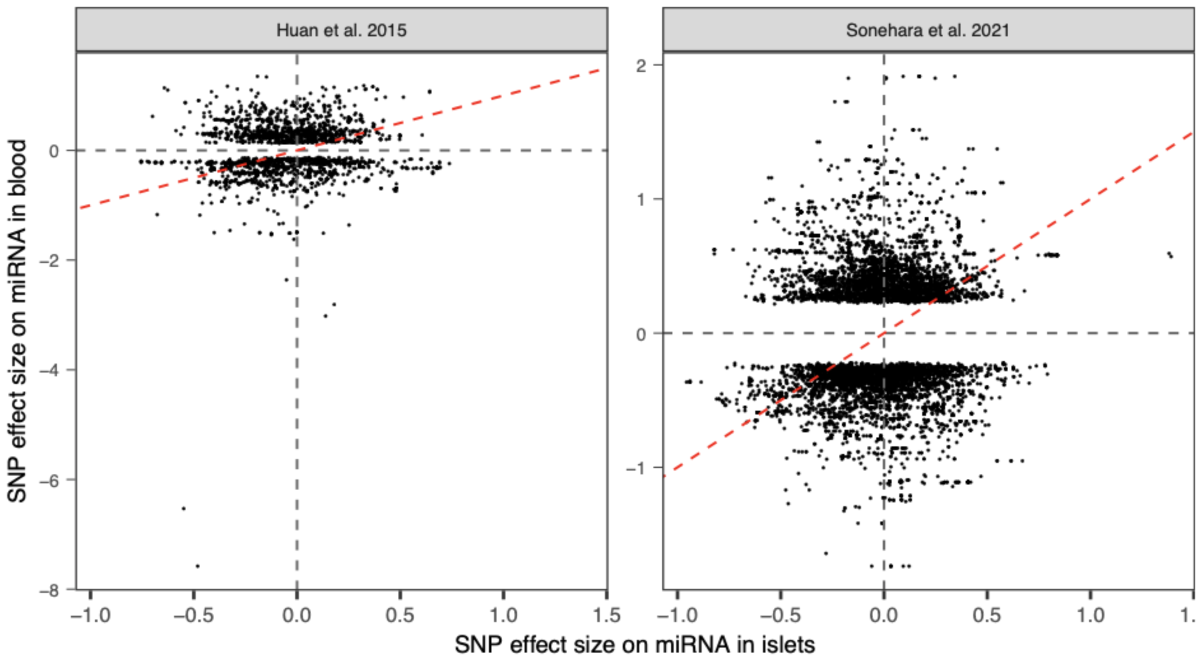


**Fig. S9. SNP-miRNA effect sizes in human pancreatic islets compared to blood.** Blood effect sizes are from published studies of genetic effects on miRNA in blood samples (Huan et al., 2015; Sonehara et al., 2021). Huan et al. released summary statistics for SNP-miRNA pairs at FDR≤10%. Sonehara et al. released summary statistics for SNP-miRNA pairs at $P \leq 0.05$. Red dashed line represents the identity line (slope=1 and intercept=0). Note: some effects are not shown due to different tag SNPs between studies even though the tag SNPs are in high linkage disequilibrium (e.g., the miR-1908 islet eQTL is tagged by rs174559 whereas the blood eQTL is tagged by rs174578 in Sonehara et al.).

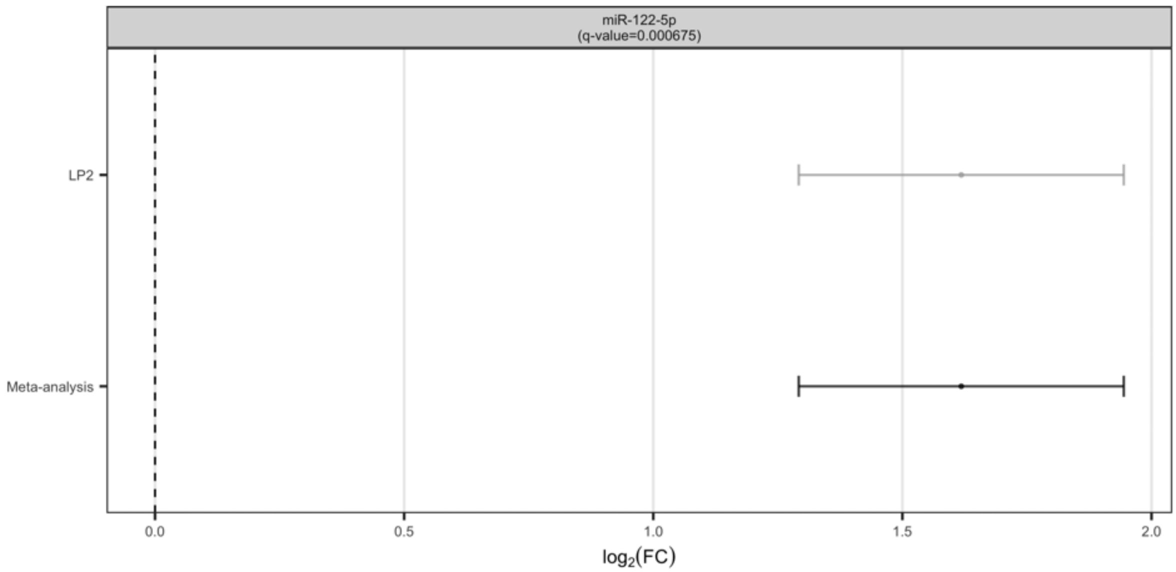


**Fig. S10. BMI-associated miRNAs.** Forest plots showing the effect of BMI-associated miRNAs ($|log_2(FC)| \geq 1$ and FDR≤5%) across the analyzed studies and meta-analysis. No point indicates that a transcript was not quantified in a specific study after processing and quality control filters. FC stands for fold change.

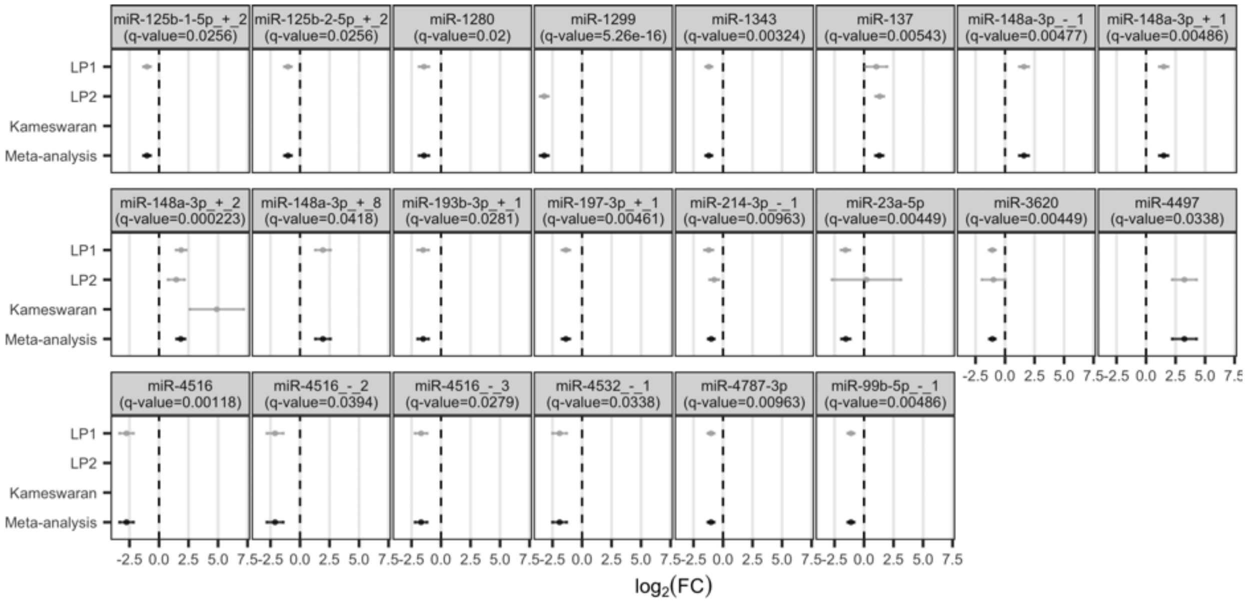


**Fig. S11. Sex-associated miRNAs.** Forest plots showing the effect of sex-associated miRNAs (male vs. female; $|log_2(FC)| \geq 1$ and FDR≤5%) across the analyzed studies and meta-analysis. No point indicates that a transcript was not quantified in a specific study after processing and quality control filters. FC stands for fold change.

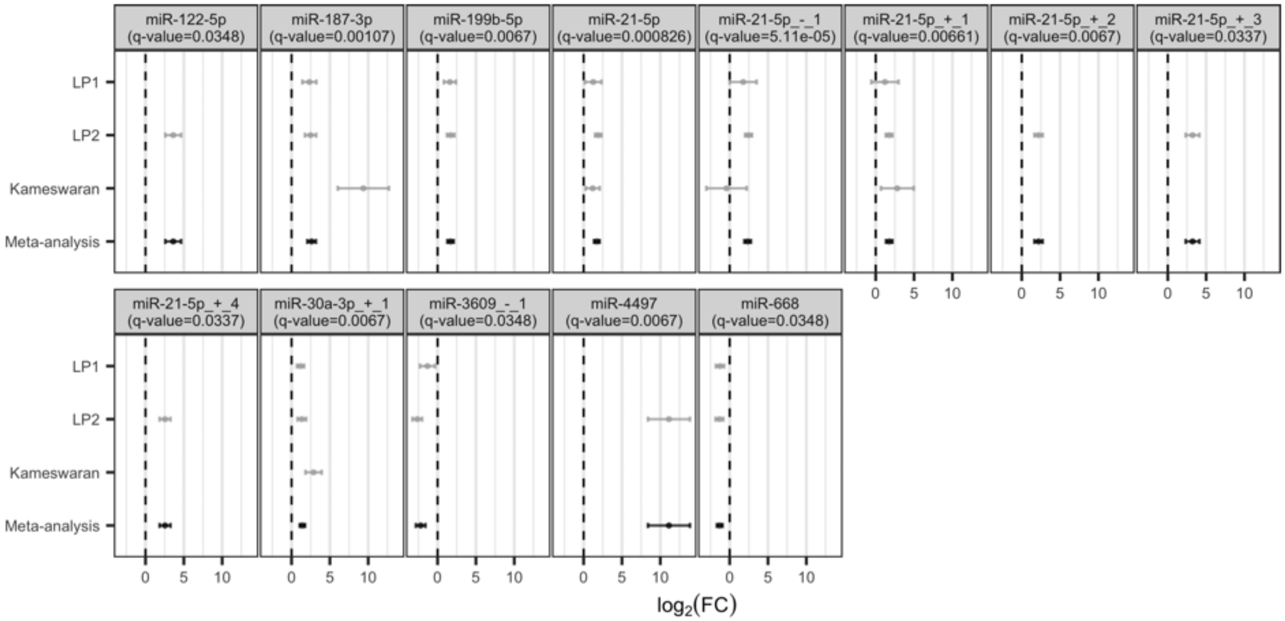


**Fig. S12. T2D-associated miRNAs.** Forest plots showing the effect of T2D-associated miRNAs ($|log_2(FC)| \geq 1$ and FDR≤5%) across the analyzed studies and meta-analysis. No point indicates that a transcript was not quantified in a specific study after processing and quality control filters. FC stands for fold change.

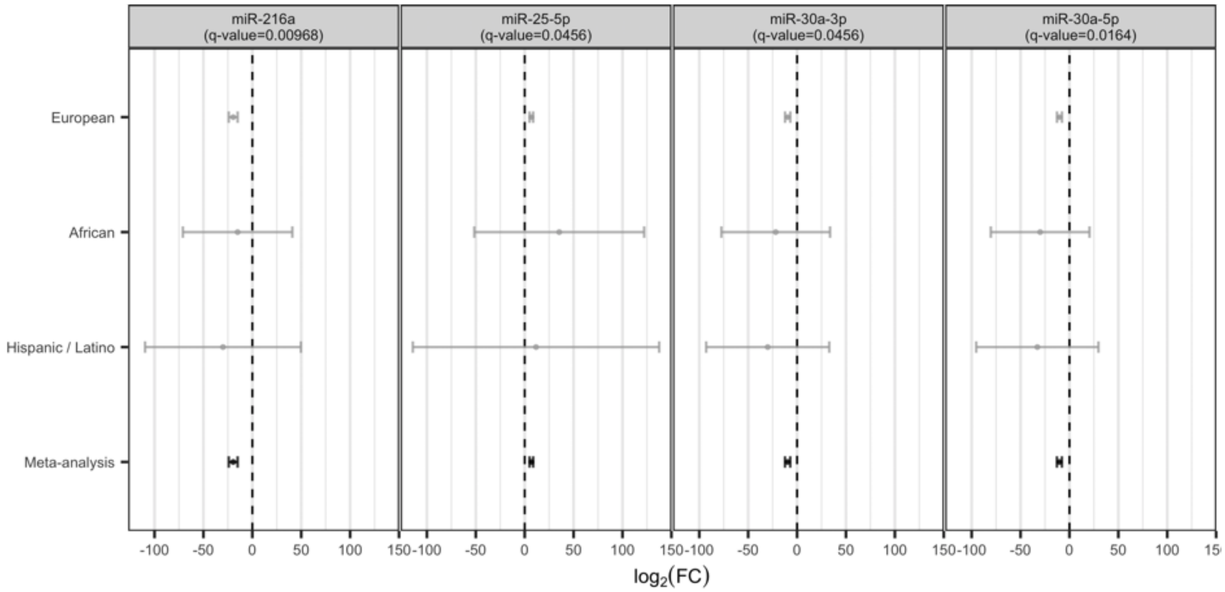


**Fig. S13. HbA1c PGS-associated miRNAs.** Forest plots showing the effect of HbA1c PGS-associated miRNAs ($|log_2(FC)| \geq 1$ and FDR≤5%) across the analyzed studies and meta-analysis. No point indicates that a transcript was not quantified in a specific study after processing and quality control filters. FC stands for fold change.

**(A)** miR-532

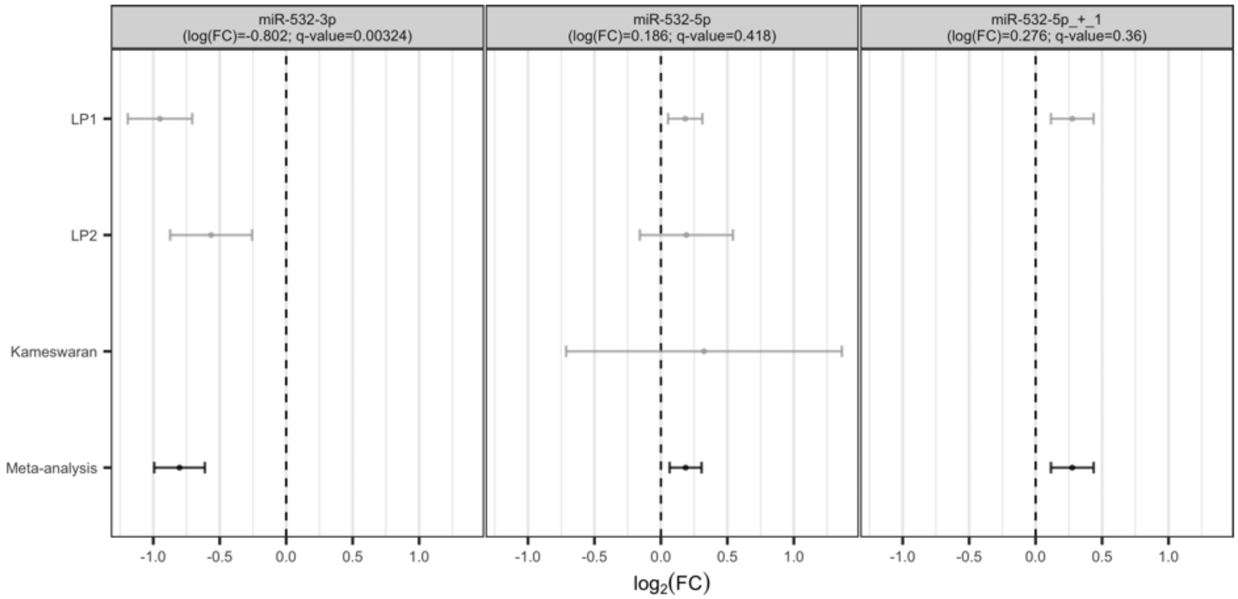


**(B)** miR-660

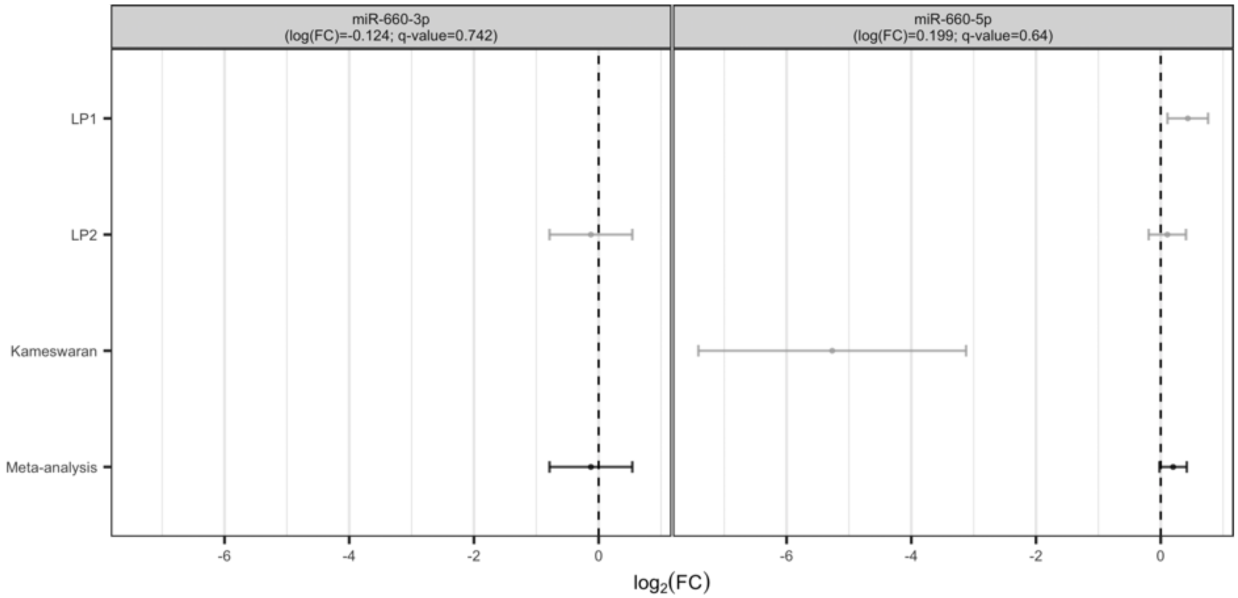


**Fig. S14. Previously reported sex-associated miRNAs.** Forest plots showing the effect of previously reported sex-associated miRNAs (male vs. female; Hall et al., 2014) in this study. We report associations with $|log_2(FC)| \geq 1$ and FDR$\leq$5% in the main text. No point indicates that a transcript was not quantified in a specific study after processing and quality control filters. FC stands for fold change.

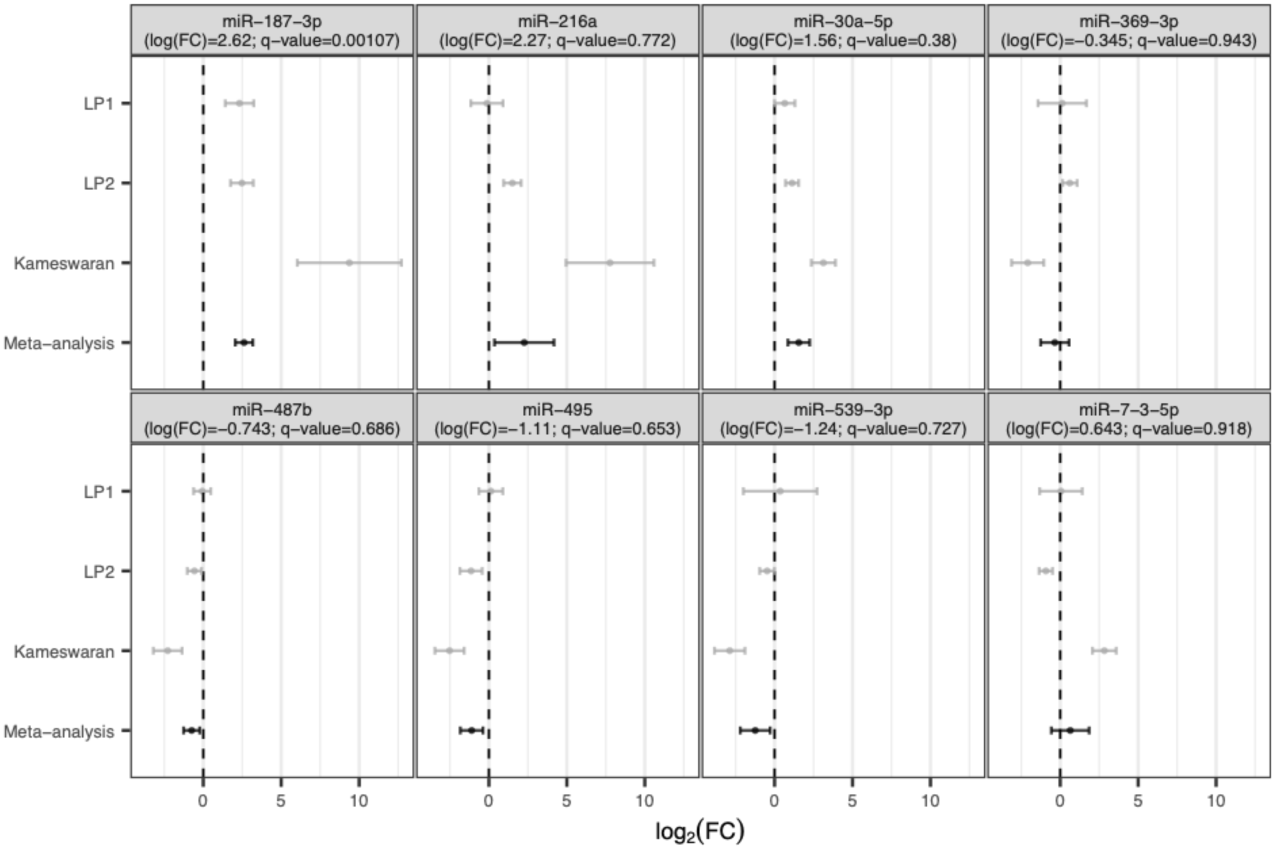


**Fig. S15. Previously reported T2D-associated miRNAs.** Forest plots showing the effect of previously reported T2D-associated miRNAs (Kameswaran et al., 2014) in this study. We report associations with $|log_2(FC)| \geq 1$ and FDR≤5% in the main text. FC stands for fold change.

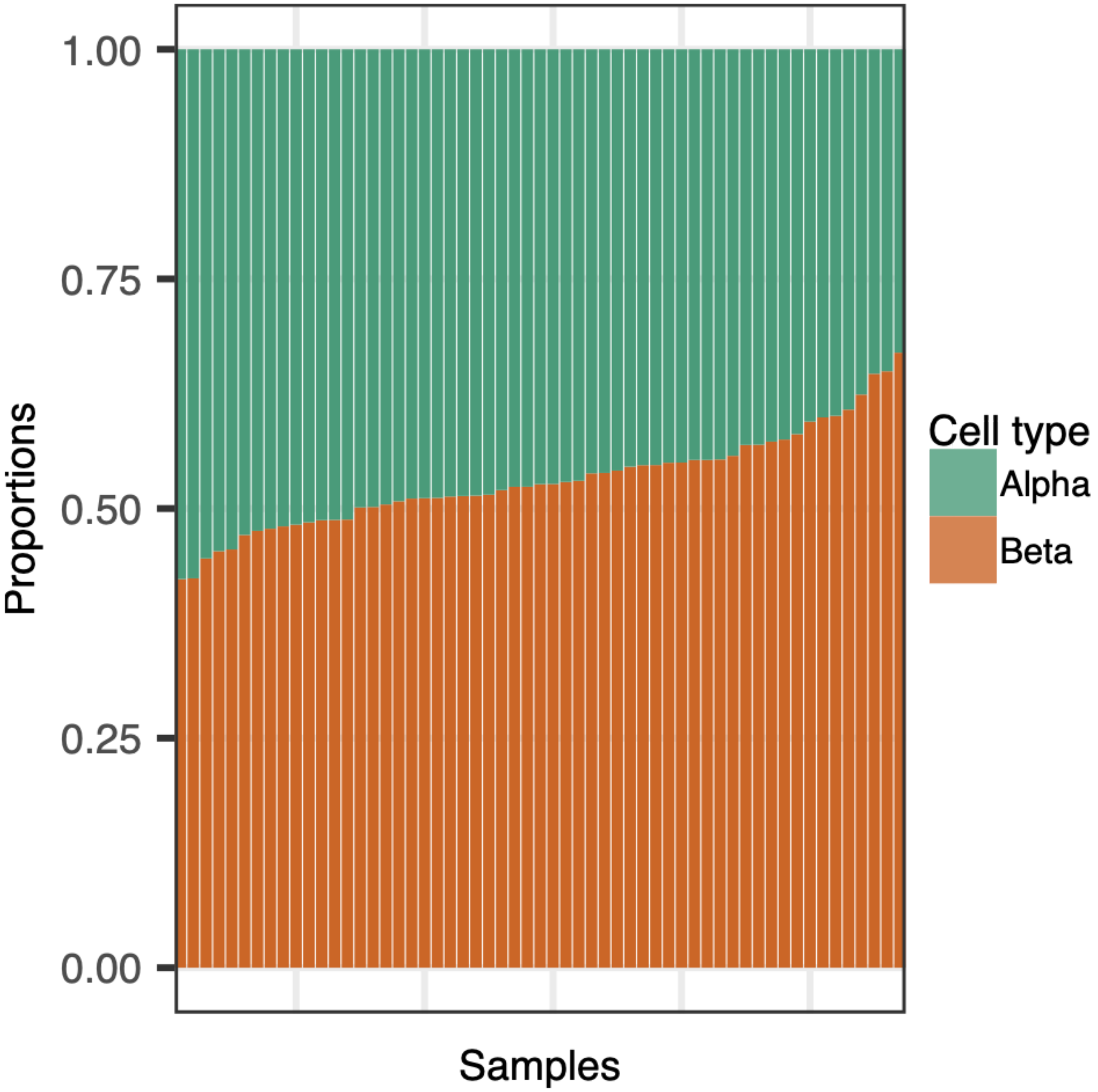


**Fig. S16. Estimated cell type proportions.** Proportion of beta and alpha cells (y-axis) in each sample (x-axis) estimated using sorted beta and alpha cells as reference (Kameswaran et al., 2014).

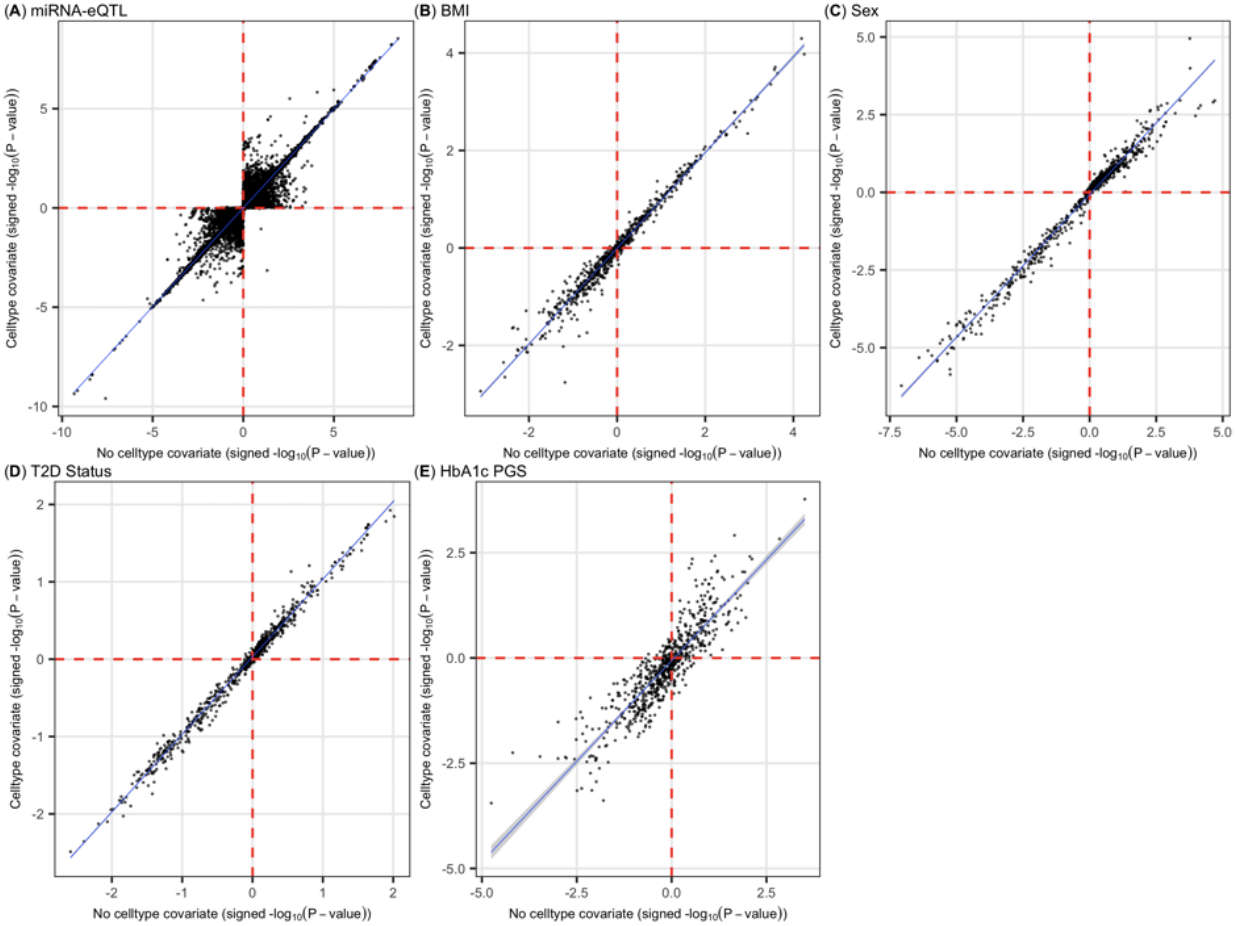


**Fig. S17. Effect of cell type proportion covariate.** Comparison of signed $-log_{10}$(*P*-values) from models with (y-axis) and without (x-axis) cell type proportions as a covariate. (A) miRNA-eQTL results. (B-E) Differential miRNA expression results across (B) BMI, (C) sex (male vs. female), (D) T2D status, and (E) HbA1c PGS.

### Supplementary Tables

**Table S1. Study characteristics.**

| | miRNA | miRNA+Genotypes | mRNA+Genotypes | miRNA+mRNA+Genotypes |
|---|---|---|---|---|
| N | 63 | 57 | 40 | 34 |
| Age (mean (SD)) | 46.38 (10.69) | 46.25 (10.48) | 45.35 (9.01) | 45.59 (8.67) |
| BMI (mean (SD)) | 28.01 (4.75) | 28.25 (4.82) | 27.8 (4.73) | 28.77 (4.11) |
| Weight (kg.) (mean (SD)) | 83.36 (17.62) | 84 (17.66) | 83.55 (16.12) | 86.13 (15.74) |
| Height (in.) (mean (SD)) | 67.96 (3.98) | 67.74 (3.85) | 67.34 (3.6) | 67.61 (3.75) |
| Sex | | | | |
| Male | 37 | 33 | 22 | 22 |
| Female | 26 | 24 | 18 | 12 |
| T2D status | | | | |
| NGT | 59 | 55 | 40 | 34 |
| T2D | 4 | 2 | 0 | 0 |

**Table S2. Characteristics of individuals in T2D status differential miRNA expression analysis.**

| | NGT | T2D |
|---|---|---|
| N | 59 | 4 |
| Age (mean (SD)) | 46.08 (10.31) | 50.75 (16.82) |
| BMI (mean (SD)) | 27.82 (4.79) | 30.52 (3.76) |
| Weight (kg.) (mean (SD)) | 82.89 (18.08) | 89.97 (6.71) |
| Height (in.) (mean (SD)) | 67.97 (4.08) | 67.75 (2.5) |
| Sex | | |
| Male | 34 | 3 |
| Female | 25 | 1 |

**Table S3. miRNA-eQTL results.** We identify 5 miRNA-eQTLs (FDR$\leq$5%). Note, empirical *P*-values were limited by the number of permutations performed in QTL mapping (max 10,000 permutations).

| miRNA | Lead SNP | Beta | Std. error | P-value | Empirical P-value |
|---|---|---|---|---|---|
| miR-3615 | *rs8064299* | -0.966411 | 0.129553 | $4.63 \times 10^{-10}$ | $1.00 \times 10^{-4}$ |
| miR-641 | *rs3730055* | 1.52844 | 0.244736 | $5.00 \times 10^{-8}$ | $1.00 \times 10^{-4}$ |
| miR-671-3p | *rs7805967* | -0.968849 | 0.150761 | $2.50 \times 10^{-8}$ | $1.00 \times 10^{-4}$ |
| miR-1343 | *rs2986402* | 0.95516 | 0.157905 | $1.06 \times 10^{-7}$ | $1.00 \times 10^{-4}$ |
| miR-1908 | *rs174559* | 0.879154 | 0.12583 | $2.88 \times 10^{-9}$ | $1.00 \times 10^{-4}$ |

**Table S4. T2D-related credible set SNPs within miRNA target sites.** SNPs from 99% credible sets for T2D and glycemic traits that lie within target sites of highly expressed miRNAs in human pancreatic islets.

| miRNA | SNP rsid | Target gene | Trait |
|---|---|---|---|
| miR-125a-5p | rs535165413 | *SLC13A3* | T2D |
| miR-301a-3p | rs145512944 | *ANKFY1* | T2D |
| miR-454-3p | rs145512944 | *ANKFY1* | T2D |
| miR-130b-3p | rs145512944 | *ANKFY1* | T2D |
| miR-130a-3p | rs145512944 | *ANKFY1* | T2D |
| miR-136-5p | rs28399585 | *CCNE2* | T2D |
| miR-149-5p | rs11655020 | *ERN1* | T2D |
| miR-182-5p | rs540515957 | *OGDH* | T2D |
| miR-186-5p | rs184830298 | *UBE2B* | T2D |
| miR-19a-3p | rs779689846 | *VPS4B* | T2D |
| miR-204-5p | rs150987217 | *PCGF3* | T2D |
| miR-212-5p | rs145527012 | *RNF10* | T2D |
| miR-330-5p | rs147493553 | *MRFAP1* | T2D |
| miR-326 | rs147493553 | *MRFAP1* | T2D |
| miR-335-5p | rs2849383 | *KDSR* | T2D |
| miR-34a-5p | rs551080524 | *HNF4A* | T2D |
| miR-369-3p | rs193171458 | *GABPB2* | T2D |
| miR-374a-5p | rs193171458 | *GABPB2* | T2D |
| miR-374b-5p | rs193171458 | *GABPB2* | T2D |
| miR-505-3p | rs34611001 | *EVC* | T2D |
| miR-543 | rs773055626 | *PPP2R2C* | T2D |
| miR-665 | rs139252054 | *TMEM41A* | T2D |
| miR-1271-5p | rs540515957 | *OGDH* | T2D |
| miR-96-5p | rs540515957 | *OGDH* | T2D |
| miR-125a-5p | rs7522956 | *WDR26* | HbA1c |
| miR-199b-3p | rs7295294 | *NAA25* | HbA1c |
| miR-200c-3p | rs3088285 | *ATXN1L* | HbA1c |
| miR-429 | rs3088285 | *ATXN1L* | HbA1c |
| miR-200b-3p | rs3088285 | *ATXN1L* | HbA1c |
| miR-200c-3p | 17:77767816_CT_C | *CBX8* | HbA1c |
| miR-429 | 17:77767816_CT_C | *CBX8* | HbA1c |
| miR-200b-3p | 17:77767816_CT_C | *CBX8* | HbA1c |
| miR-339-5p | rs144010860 | *PLD1* | HbA1c |
| miR-339-5p | rs1076504 | *PLD1* | HbA1c |
| miR-485-5p | rs9605070 | *ZDHHC8* | HbA1c |
| miR-532-3p | rs1464569 | *NICN1* | HbA1c |
| miR-146a-5p | rs72763690 | *LUC7L* | Blood glucose |

**Table S5. T2D status, age, sex, and BMI differential expression covariates.** Covariates accounted for within each miRNA library preparation for differential miRNA expression analyses across T2D status, age, sex, and BMI.

| | Library preparation | | |
|---|---|---|---|
| Tested trait | LP1 | LP2 | Kameswaran |
| T2D status | *sex, age, RUVseq Factors 1 & 2* | *sex, RUVseq Factors 1 & 2* | *sex, age, RUVseq Factors 1 & 2* |
| Sex | *T2D status, age, RUVseq Factors 1 & 2* | *T2D status, RUVseq Factors 1 & 2* | *T2D status, age, RUVseq Factors 1 & 2* |
| Age | *T2D status, sex, RUVseq Factors 1 & 2* | *T2D status, sex, RUVseq Factor 1* | *T2D status, sex, RUVseq Factors 1 & 2* |
| BMI | *T2D status, sex, age, RUVseq Factors 1 & 2* | *T2D status, sex, RUVseq Factor 1* | *T2D status, sex, RUVseq Factors 1 & 2* |

**Table S6. PGS differential expression covariates.** Covariates accounted for within each ancestry for the PGS differential miRNA expression analyses.

| | Ancestry | | |
|---|---|---|---|
| Tested trait | European | African | Hispanic/Latino |
| PGS | *sex, age, genetic PCs 1-4, RUVseq Factors 1 & 2* | *sex, age, RUVseq Factors 1 & 2* | *sex, RUVseq Factors 1 & 2* |

#### SI References


1. M. C. Gershengorn, *et al.*, Epithelial-to-mesenchymal transition generates proliferative human islet precursor cells. *Science* **306**, 2261–2264 (2004).
2. N. Lawlor, *et al.*, Multiomic Profiling Identifies cis-Regulatory Networks Underlying Human Pancreatic β Cell Identity and Function. *Cell Rep.* **26**, 788-801.e6 (2019).
3. A. Manichaikul, *et al.*, Robust relationship inference in genome-wide association studies. *Bioinformatics* **26**, 2867–2873 (2010).
4. J. Novembre, *et al.*, Genes mirror geography within Europe. *Nature* **456**, 98–101 (2008).
5. D. Taliun, *et al.*, LASER server: ancestry tracing with genotypes or sequence reads. *Bioinformatics* **33**, 2056–2058 (2017).
6. A. L. Price, *et al.*, Long-range LD can confound genome scans in admixed populations. *Am. J. Hum. Genet.* **83**, 132–135 (2008).
7. M. E. Weale, Quality control for genome-wide association studies. *Methods Mol. Biol.* **628**, 341–372 (2010).
8. A. L. Price, *et al.*, Principal components analysis corrects for stratification in genome-wide association studies. *Nat. Genet.* **38**, 904–909 (2006).
9. N. Patterson, A. L. Price, D. Reich, Population structure and eigenanalysis. *PLoS Genet.* **2**, e190 (2006).
10. C. A. Tracy, H. Widom, Level spacing distributions and the Bessel kernel. *Commun.Math. Phys.* **161**, 289–309 (1994).
11. K. W. Currin, *et al.*, Genetic effects on liver chromatin accessibility identify disease regulatory variants. *Am. J. Hum. Genet.* **108**, 1169–1189 (2021).
12. S. Das, *et al.*, Next-generation genotype imputation service and methods. *Nat. Genet.* **48**, 1284–1287 (2016).
13. P.-R. Loh, P. F. Palamara, A. L. Price, Fast and accurate long-range phasing in a UK Biobank cohort. *Nat. Genet.* **48**, 811–816 (2016).
14. O. Delaneau, J. Marchini, J.-F. Zagury, A linear complexity phasing method for thousands of genomes. *Nat. Methods* **9**, 179–181 (2011).
15. S. McCarthy, *et al.*, A reference panel of 64,976 haplotypes for genotype imputation. *Nat. Genet.* **48**, 1279–1283 (2016).
16. M. Kanke, J. Baran-Gale, J. Villanueva, P. Sethupathy, miRquant 2.0: an Expanded Tool for Accurate Annotation and Quantification of MicroRNAs and their isomiRs from Small RNA-Sequencing Data. *J. Integr. Bioinform.* **13**, 307 (2016).
17. M. Martin, Cutadapt removes adapter sequences from high-throughput sequencing reads. *EMBnet j.* **17**, 10 (2011).
18. B. Langmead, C. Trapnell, M. Pop, S. L. Salzberg, Ultrafast and memory-efficient alignment of short DNA sequences to the human genome. *Genome Biol.* **10**, R25 (2009).
19. M. David, M. Dzamba, D. Lister, L. Ilie, M. Brudno, SHRiMP2: sensitive yet practical SHort Read Mapping. *Bioinformatics* **27**, 1011–1012 (2011).
20. G. Jun, *et al.*, Detecting and estimating contamination of human DNA samples in sequencing and array-based genotype data. *Am. J. Hum. Genet.* **91**, 839–848 (2012).

21. J. Rozowsky, *et al.*, exceRpt: A Comprehensive Analytic Platform for Extracellular RNA Profiling. *Cell Syst.* **8**, 352-357.e3 (2019).

22. A. Dobin, *et al.*, STAR: ultrafast universal RNA-seq aligner. *Bioinformatics* **29**, 15–21 (2013).

23. S. W. Hartley, J. C. Mullikin, QoRTs: a comprehensive toolset for quality control and data processing of RNA-Seq experiments. *BMC Bioinformatics* **16**, 224 (2015).

24. C. Lippert, F. P. Casale, B. Rakitsch, O. Stegle, LIMIX: genetic analysis of multiple traits. *BioRxiv* (2014) https:/doi.org/10.1101/003905.

25. O. Stegle, L. Parts, M. Piipari, J. Winn, R. Durbin, Using probabilistic estimation of expression residuals (PEER) to obtain increased power and interpretability of gene expression analyses. *Nat. Protoc.* **7**, 500–507 (2012).

26. L. Chen, *et al.*, Genetic drivers of epigenetic and transcriptional variation in human immune cells. *Cell* **167**, 1398-1414.e24 (2016).

27. L. J. Scott, *et al.*, The genetic regulatory signature of type 2 diabetes in human skeletal muscle. *Nat. Commun.* **7**, 11764 (2016).

28. G. E. Hoffman, Correcting for population structure and kinship using the linear mixed model: theory and extensions. *PLoS ONE* **8**, e75707 (2013).

29. H. Ongen, A. Buil, A. A. Brown, E. T. Dermitzakis, O. Delaneau, Fast and efficient QTL mapper for thousands of molecular phenotypes. *Bioinformatics* **32**, 1479–1485 (2016).

30. J. D. Storey, R. Tibshirani, Statistical significance for genomewide studies. *Proc Natl Acad Sci USA* **100**, 9440–9445 (2003).

31. C. Giambartolomei, *et al.*, Bayesian test for colocalisation between pairs of genetic association studies using summary statistics. *PLoS Genet.* **10**, e1004383 (2014).

32. H. Guo, *et al.*, Integration of disease association and eQTL data using a Bayesian colocalisation approach highlights six candidate causal genes in immune-mediated diseases. *Hum. Mol. Genet.* **24**, 3305–3313 (2015).

33. J. Chen, *et al.*, The trans-ancestral genomic architecture of glycemic traits. *Nat. Genet.* **53**, 840–860 (2021).

34. N. Sinnott-Armstrong, *et al.*, Genetics of 35 blood and urine biomarkers in the UK Biobank. *Nat. Genet.* **53**, 185–194 (2021).

35. A. Mahajan, *et al.*, Fine-mapping type 2 diabetes loci to single-variant resolution using high-density imputation and islet-specific epigenome maps. *Nat. Genet.* **50**, 1505–1513 (2018).

36. Pan-UKB team, Pan-UK Biobank. *https://pan.ukbb.broadinstitute.org* (2020) (March 18, 2021).

37. A. Viñuela, *et al.*, Genetic variant effects on gene expression in human pancreatic islets and their implications for T2D. *Nat. Commun.* **11**, 4912 (2020).

38. J. Millstein, G. K. Chen, C. V. Breton, cit: hypothesis testing software for mediation analysis in genomic applications. *Bioinformatics* **32**, 2364–2365 (2016).

39. D. L. Taylor, *et al.*, Integrative analysis of gene expression, DNA methylation, physiological traits, and genetic variation in human skeletal muscle. *Proc Natl Acad Sci USA* **116**, 10883–10888 (2019).

40. T. Huan, *et al.*, Genome-wide identification of microRNA expression quantitative trait loci. *Nat. Commun.* **6**, 6601 (2015).

41. K. Sonehara, *et al.*, Genetic architecture of microRNA expression and its link to complex diseases in the Japanese population. *Hum. Mol. Genet.* (2021) https:/doi.org/10.1093/hmg/ddab361.

42. W. W. Greenwald, *et al.*, Pancreatic islet chromatin accessibility and conformation reveals distal enhancer networks of type 2 diabetes risk. *Nat. Commun.* **10**, 2078 (2019).

43. H. Li, R. Durbin, Fast and accurate long-read alignment with Burrows-Wheeler transform. *Bioinformatics* **26**, 589–595 (2010).

44. B. van de Geijn, G. McVicker, Y. Gilad, J. K. Pritchard, WASP: allele-specific software for robust molecular quantitative trait locus discovery. *Nat. Methods* **12**, 1061–1063 (2015).

45. Y. Benjamini, Y. Hochberg, Controlling the false discovery rate: a practical and powerful approach to multiple testing. *Journal of the Royal Statistical Society: Series B (Methodological)* **57**, 289–300 (1995).

46. A. R. Quinlan, I. M. Hall, BEDTools: a flexible suite of utilities for comparing genomic features. *Bioinformatics* **26**, 841–842 (2010).

47. V. Agarwal, G. W. Bell, J.-W. Nam, D. P. Bartel, Predicting effective microRNA target sites in mammalian mRNAs. *eLife* **4** (2015).

48. S. Fairley, E. Lowy-Gallego, E. Perry, P. Flicek, The International Genome Sample Resource (IGSR) collection of open human genomic variation resources. *Nucleic Acids Res.* **48**, D941–D947 (2020).

49. K. L. Howe, *et al.*, Ensembl 2021. *Nucleic Acids Res.* **49**, D884–D891 (2021).

50. A. Kozomara, M. Birgaoanu, S. Griffiths-Jones, miRBase: from microRNA sequences to function. *Nucleic Acids Res.* **47**, D155–D162 (2019).

51. A. Kozomara, S. Griffiths-Jones, miRBase: annotating high confidence microRNAs using deep sequencing data. *Nucleic Acids Res.* **42**, D68-73 (2014).

52. A. Kozomara, S. Griffiths-Jones, miRBase: integrating microRNA annotation and deep-sequencing data. *Nucleic Acids Res.* **39**, D152-7 (2011).

53. S. Griffiths-Jones, H. K. Saini, S. van Dongen, A. J. Enright, miRBase: tools for microRNA genomics. *Nucleic Acids Res.* **36**, D154-8 (2008).

54. S. Griffiths-Jones, R. J. Grocock, S. van Dongen, A. Bateman, A. J. Enright, miRBase: microRNA sequences, targets and gene nomenclature. *Nucleic Acids Res.* **34**, D140-4 (2006).

55. S. Griffiths-Jones, The microRNA Registry. *Nucleic Acids Res.* **32**, D109-11 (2004).

56. S. Purcell, *et al.*, PLINK: a tool set for whole-genome association and population-based linkage analyses. *Am. J. Hum. Genet.* **81**, 559–575 (2007).

57. S. Purcell, PLINK v1.07 (August 18, 2021).

58. Y. Tanigawa, N. Sinnott-Armstrong, M. Rivas, The snpnet polygenic risk score coefficients for 35 lab biomarkers described in “Genetics of 35 blood and urine biomarkers in the UK Biobank.” *The NIH Figshare Archive* (2020) https:/doi.org/10.35092/yhjc.12298838.v1.

59. T. Ge, C.-Y. Chen, Y. Ni, Y.-C. A. Feng, J. W. Smoller, Polygenic prediction via Bayesian regression and continuous shrinkage priors. *Nat. Commun.* **10**, 1776 (2019).

60. V. Kameswaran, *et al.*, Epigenetic regulation of the DLK1-MEG3 microRNA cluster in human type 2 diabetic islets. *Cell Metab.* **19**, 135–145 (2014).

61. A. R. Martin, *et al.*, Human Demographic History Impacts Genetic Risk Prediction across Diverse Populations. *Am. J. Hum. Genet.* **100**, 635–649 (2017).

62. M. I. Love, W. Huber, S. Anders, Moderated estimation of fold change and dispersion for RNA-seq data with DESeq2. *Genome Biol.* **15**, 550 (2014).

63. D. Risso, J. Ngai, T. P. Speed, S. Dudoit, Normalization of RNA-seq data using factor analysis of control

genes or samples. *Nat. Biotechnol.* **32**, 896–902 (2014).

64. C. Prada, D. Lima, H. Nakaya, *MetaVolcanoR: Gene Expression Meta-analysis Visualization Tool* (R package, 2021).

65. W. G. Cochran, The Combination of Estimates from Different Experiments. *Biometrics* **10**, 101 (1954).
